# Physiological mapping and prediction reveal stress-responsive translation initiation landscapes

**DOI:** 10.64898/2026.11.03.763266

**Authors:** Kazuya Ichihara, Taichi Shiraishi, Haruhiko Morita, Chisa Shiraishi, Mina Hirata, Naoto Yamashiki, Selena Setsu, Kotaro Tsuboyama, Atsushi Hatano, Serina Tokita, Emi Mishiro-Sato, Ayako Sakamoto, Takuhiro Ito, Kazumi Abe, Masahide Seki, Yutaka Suzuki, Masahiko Ajiro, Akihide Yoshimi, Takayuki Kanaseki, Masaki Matsumoto, Shungo Adachi, Pavel V. Baranov, Teppei Shimamura, Akinobu Matsumoto

## Abstract

Translation initiation defines the protein output of the transcriptome, but its physiological dynamics in cells and tissues remain incompletely characterized. Here we develop TISCA2, an inhibitor-free method that captures eIF5B-associated initiating ribosomes to map translation initiation sites genome-wide. By directly identifying start sites rather than inferring translation from elongating-ribosome periodicity, TISCA2 improves the detection of short and overlapping translons. Productive translation of a subset of TISCA2-identified translons is supported by N-terminal proteomics and HLA-I immunopeptidomics. The arsenite-induced integrated stress response broadly redistributes start-site usage, reducing upstream initiation in favor of downstream initiation. TISCA2 further reveals tissue-specific initiation landscapes in vivo, with increased detection of downstream candidate start sites in the testis. Finally, we develop IncepTIS, a sequence-based deep learning model that predicts position-resolved and condition-dependent initiation landscapes. IncepTIS-guided optimization generates de novo mRNA 5′ leaders with stress-responsive initiation patterns. Together, TISCA2 and IncepTIS provide an experimental and computational framework for mapping, predicting and engineering translation initiation in mammalian cells and tissues.

## Introduction

Translation initiation determines where ribosomes begin decoding mRNAs and thereby defines the protein output of the transcriptome^1–3^. Although eukaryotic translation has long been viewed through the lens of annotated, predominantly AUG-initiated coding sequences (CDSs), ribosome profiling and related approaches have revealed a broader translational landscape^4–6^. Ribosomes can initiate both upstream and downstream of annotated CDS start sites. Translation initiation also occurs at near-cognate start codons such as CUG, GUG, ACG and UUG, generally less efficiently than at AUG codons^7,8^. Ribosome profiling has revealed that such non-AUG initiation is widespread and may lead to the synthesis of alternative N-terminal proteoforms, some of which are under evolutionary selection^5,9–13^.

This expanding translation landscape has prompted efforts to standardize the annotation of translation beyond regions classified as protein-coding by current annotation pipelines^14^. The TransCODE consortium produces and maintains a catalogue of translated regions (or translons^14^) that are supported purely by ribosome profiling data^15,16^. Some products of these translons are stable, can be detected with proteomics approaches and are biologically active, whereas others are unstable or remain undetected, and their biological roles remain unclear^17–19^.

Despite these advances, important blind spots remain. Many approaches for detecting translation rely on the reading-frame bias reflected in Ribo-seq triplet periodicity and on the distribution of elongating-ribosome footprints across a translon^20–23^. Although approaches for detecting translation in overlapping reading frames exist, detection accuracy depends strongly on read coverage and the relative translation levels of the overlapping translons^24,25^. Moreover, even when translation is detected, precise start-codon assignment remains challenging, particularly for near-cognate or condition-dependent initiation, or when multiple start codons are used within the same open reading frame (ORF). This is known to occur when translation initiation is downstream of annotated AUGs or occurs at multiple weak non-AUG starts^26,27^. Translons initiated at different sites within the same ORF share the same reading frame and therefore cannot be distinguished by triplet periodicity alone in their overlapping regions. Moreover, changes in footprint density associated with additional initiation can be confounded by variation in elongation rates, including ribosome pausing.

Translation initiation site (TIS) mapping has addressed some of these limitations using inhibitors such as harringtonine or lactimidomycin to enrich ribosomes near start codons^5,9,10^. However, inhibitor-based profiling can yield initiation-like signals that may reflect the arrest of elongating ribosomes rather than genuine translation initiation^28^. We previously developed TISCA, which integrates selective translation complex profiling sequencing (Sel-TCP-seq) with global translation initiation sequencing (GTI-seq) and Ribo-seq to improve TIS detection^29^, but its dependence on inhibitor-based data leaves these artifacts unresolved. In vivo, delayed and tissue-dependent drug penetration further complicates precise start-site mapping, particularly in barrier-protected organs such as brain and testis^30,31^. An inhibitor-free approach is therefore needed to define physiological initiation landscapes in cells and tissues.

Start-site selection is also dynamically regulated by cellular state. For example, during the integrated stress response (ISR), eIF2α phosphorylation suppresses global translation while enabling selective translation of stress-regulated mRNAs such as *ATF4* through upstream ORF (uORF)-dependent mechanisms^32^. However, studies of stress-regulated initiation have largely focused on annotated 5′ untranslated regions (UTRs) and the balance between uORFs and CDSs^33–36^. Whether stress broadly redistributes initiation across entire transcripts, including CDSs and annotated 3′ UTRs, remains largely unknown.

A model that predicts TISs from sequence and cellular state would allow condition-dependent changes in initiation to be estimated for transcripts and sequence variants that have not been profiled experimentally, enabling the prioritization of candidates for experimental testing. Several deep learning models have been developed to predict translation initiation sites or translational output more broadly, either from sequence alone or from sequence together with RNA expression data^37–40^. However, these models were not designed to predict stress-dependent changes in start-site usage for a given transcript. Moreover, models trained on annotated start sites may be biased toward known initiation events, highlighting the value of condition-resolved experimental initiation profiles as training data.

Here we develop TISCA2, an inhibitor-free method that captures eIF5B-associated initiating ribosomes to map TISs genome-wide. By identifying start sites without relying on the triplet periodicity of Ribo-seq footprints, TISCA2 resolves short and overlapping translons, including internal ORFs (intORFs) and truncated variant translons (Fig. 1) that are difficult to capture by conventional Ribo-seq-based approaches. TISCA2 reveals that the arsenite-induced ISR broadly redistributes translation initiation toward downstream sites within CDSs and annotated 3′ UTRs and further uncovers tissue-specific initiation landscapes in vivo. Finally, we develop IncepTIS, a sequence-based deep learning model that predicts position-resolved, condition-dependent initiation landscapes and enables the prospective design of stress-responsive translational elements. Together, TISCA2 and IncepTIS provide a framework for mapping, predicting and engineering translation initiation in mammalian cells and tissues.

**Fig. 1:**
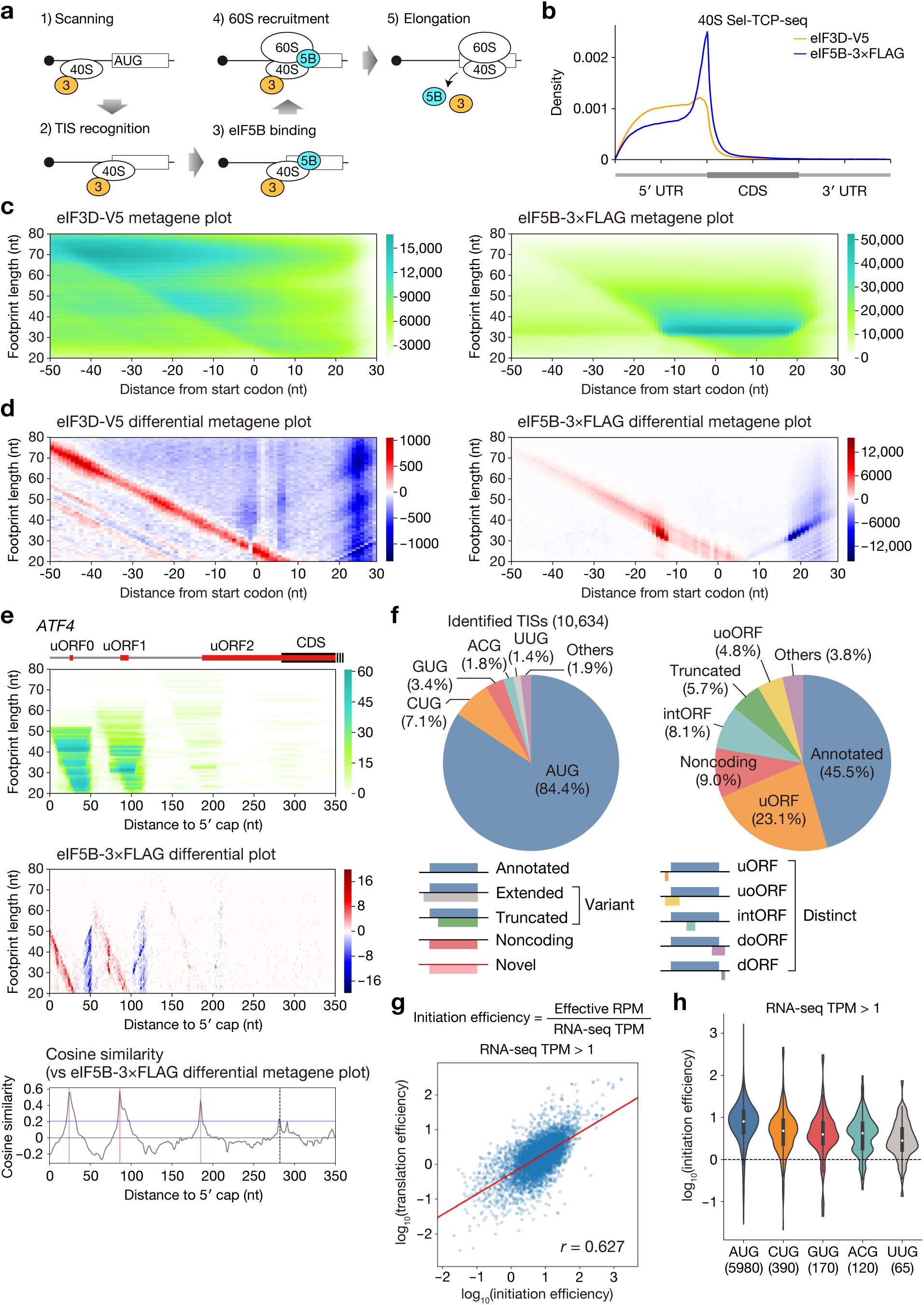
eIF5B-associated ribosome footprints enable inhibitor-free identification and quantification of translation initiation sites. **a**, Schematic of translation initiation and the association of eIF3 and eIF5B with sequential initiation intermediates. **b**, Metagene profiles of eIF3D-V5 and eIF5B- 3×FLAG Sel-TCP-seq footprints in HEK293T cells. Representative data of two biological replicates are shown. All reads were mapped to all human protein-coding transcripts with a 5′-UTR and 3′-UTR of >100 nt (*n* = 52,692). **c**, Two-dimensional metagene distributions of eIF3D-V5 and eIF5B-3×FLAG footprints around annotated start codons. Footprint length is plotted against the distance from the annotated start codon. Representative data of two biological replicates are shown. All reads were mapped to protein-coding transcripts expressed in HEK293T cells with a 5′-UTR of >100 nt (*n* = 18,362). **d**, Differential metagene plots generated from the footprint distributions shown in **c**. **e**, Representative analysis of the *ATF4* transcript. The positions of uORF0, uORF1, uORF2 and the CDS are shown above the eIF5B- 3×FLAG footprint distribution. The differential footprint plot and cosine similarity to the eIF5B-3×FLAG differential metagene signature are shown below. Dashed vertical lines indicate the initiation sites of the indicated translons. Horizontal blue line indicates the cutoff value (0.206). **f**, Start-codon composition and translon type classification of 10,634 TISs identified by TISCA2 in HEK293T cells. The schematic below shows the types of canonical and non-canonical translons. **g**, Relationship between initiation efficiency (IE) and translation efficiency (TE). IE was calculated as TIS-proximal eIF5B effective reads per million (RPM) normalized to RNA-seq transcripts per million (TPM). Transcripts with RNA-seq TPM > 1 were included. Red line indicates the regression curve. The Pearson’s correlation coefficient is shown. **h**, Distribution of IE for TISs initiated at the indicated AUG and near-cognate codons. Only transcripts with RNA-seq TPM > 1 were included. The number of TISs in each group is indicated below the graph.

## Results

### eIF5B-associated footprints mark translation initiation sites

Eukaryotic translation initiation proceeds through scanning of the 48S pre-initiation complex (PIC), followed by start-codon recognition and recruitment of the 60S ribosomal subunit to form an elongation-competent 80S ribosome (Fig. 1a). eIF3 is associated with scanning 40S complexes and contributes to multiple steps of the initiation process^41^, whereas eIF5B is bound to late initiation complexes after start-codon recognition and before productive elongation^42–44^. We therefore reasoned that footprints of eIF5B-associated ribosomes should be enriched at TISs, whereas eIF3D-associated footprints should more broadly report locations of scanning 40S complexes.

To test this idea, we performed Sel-TCP-seq using HEK293T knock-in cells expressing either eIF3D-V5 or eIF5B-3×FLAG (Extended Data Fig. 1a,b). The eIF5B-3×FLAG libraries were highly reproducible between biological replicates (Extended Data Fig. 1c). Transcript-level footprint abundances were also compared between eIF3D and eIF5B profiles (Extended Data Fig. 1d). Metagene analysis revealed a sharp enrichment of eIF5B-3×FLAG footprints around annotated start codons, in contrast to the broader 5′ UTR distribution of eIF3D-V5 footprints (Fig. 1b and Extended Data Fig. 1e).

When both footprint position and length were considered, eIF3D- and eIF5B-associated footprints showed distinct two-dimensional patterns around annotated TISs. Compared with eIF3D-V5, eIF5B-3×FLAG footprints contained fewer long protected fragments of more than 60 nt near start codons (Fig. 1c), consistent with staged structural changes of scanning and late-initiation 40S complexes^41^. To extract TIS-proximal features from these broad and overlapping footprint distributions, we calculated differential metagene plots by measuring position-to-position changes in the two-dimensional footprint density around annotated start codons. This edge-detection approach revealed a clearer and more stereotyped pattern for eIF5B-3×FLAG than for eIF3D-V5 (Fig. 1d), suggesting that the shape of the eIF5B-associated differential footprint could be used as a signature of translation initiation.

To detect this signature on individual transcripts, we calculated the cosine similarity between the eIF5B differential metagene pattern derived from annotated AUG start sites and the differential footprint distributions within sliding 60-nt windows along each transcript (Extended Data Fig. 1f). At *ATF4*, a transcript with multiple uORFs and stress-regulated reinitiation^45,46^, eIF5B-3×FLAG differential footprints produced discrete signals at uORF0, uORF1, uORF2 and the main ORF (Fig. 1e). Cosine similarity showed sharp peaks at each of these start sites, including the annotated *ATF4* start codon, despite the low basal translation of the ATF4 main ORF. These results indicate that eIF5B-associated footprint shape can identify both strong and weak TISs without harringtonine or lactimidomycin pretreatment.

### TISCA2 enables inhibitor-free genome-wide TIS detection and quantification of initiation efficiency

We next developed TISCA2, an inhibitor-free framework for genome-wide TIS identification based on the eIF5B-associated footprint signature. We generated an expressed transcript-isoform catalog by Nanopore long-read sequencing and considered AUG and near-cognate codons differing from AUG by one nucleotide (including AAG and AGG) as candidate TISs). Candidates were filtered by footprint-length coverage and scored by cosine similarity to the eIF5B differential metagene signature. A cutoff balancing precision and recall, followed by local-maximum filtering, was used to define TISCA2-positive sites (Extended Data Fig. 2a–e).

TISCA2 identified 10,634 TISs in HEK293T cells (Fig. 1f). Most detected TISs were AUG codons (84.4%), followed by CUG (7.1%), GUG (3.4%), ACG (1.8%) and UUG (1.4%) start codons, consistent with previous reporter-based analyses showing that near-cognate codons differ widely in initiation potential^11,47^. Many transcripts contained multiple TISs (Extended Data Fig. 2f), indicating extensive start-site heterogeneity. In contrast to the first-generation TISCA approach, which incorporated inhibitor-based initiation profiling, TISCA2 detected very few AAG or AGG TISs (Fig. 1f). Some previously reported initiation-like signals at AAG and AGG codons have been attributed to the arrest of elongating ribosomes by harringtonine or lactimidomycin^28^. The low frequency of these candidates in TISCA2 is consistent with reduced contributions from drug-induced elongation artifacts.

Classification of TISCA2-defined translons showed that close to half corresponded to annotated CDS TISs (45.5%), whereas the remaining sites mapped to other categories, including uORFs (23.1%), noncoding transcripts (9.0%), intORFs (8.1%), truncated ORFs (5.7%) and upstream overlapping ORFs (uoORFs) (4.8%) (Fig. 1f). Thus, TISCA2 captures both annotated protein-coding translons (CDSs) and newly identified translons, including those internal to or overlapping with annotated CDSs. For subsequent comparisons, we grouped extended and truncated ORFs as variant translons and uORFs, uoORFs, intORFs, doORFs and dORFs as distinct translons (Fig. 1f).

Because distributions of eIF5B-associated footprints varied in both shape and intensity, we next asked whether their signal strength could provide a quantitative measure of initiation activity. We defined initiation efficiency (IE) as effective TIS-proximal eIF5B footprint counts, expressed as reads per million (RPM), divided by RNA-seq transcript abundance, expressed as transcripts per million (TPM) (Extended Data Fig. 2g). IE correlated with translational efficiency (TE) calculated from Ribo-seq and RNA-seq data^6^ (*r* = 0.627; Fig. 1g), indicating that eIF5B occupancy provides a quantitative proxy for initiation activity. Effective read counts were correlated with cosine similarity, but read depth alone did not distinguish TIS-like from non-TIS-like footprint shapes, highlighting the importance of the shape-based TISCA2 score (Extended Data Fig. 2h). Finally, stratification of IE by start codon revealed the highest IE for AUG and lower but detectable IE for major near-cognate codons, with CUG showing the highest IE among them (Fig. 1h). Across non-AUG codon types, IE correlated with the total number of detected TISs for that codon type. Together, these results establish TISCA2 as a reliable inhibitor-free method for identifying and quantifying physiological translation initiation events.

### TISCA2 improves detection of distinct translons

We next benchmarked TISCA2 against ORF-RATER, a regression-based method that identifies translons from ribosome-profiling read distributions^22^. To make the comparison as direct as possible, we incorporated TI-seq-derived start-site information into the ORF-RATER analysis and used AUG and near-cognate codons as candidate starts. TI-seq metagene analysis confirmed the expected accumulation of ribosome footprints near annotated start codons (Extended Data Fig. 3a).

Using an ORF-RATER score cutoff of 0.8, ORF-RATER identified 7,211 translons in HEK293T cells. Of these, 75.2% initiated at AUG codons, whereas CUG, AAG, GUG and AGG accounted for 7.7%, 4.6%, 3.1% and 2.4%, respectively (Fig. 2a). ORF-RATER also showed higher fractions of AAG- and AGG-associated candidates than TISCA2 (AAG, 4.6% versus 0.3%; AGG, 2.4% versus 0.4%, respectively).

**Fig. 2:**
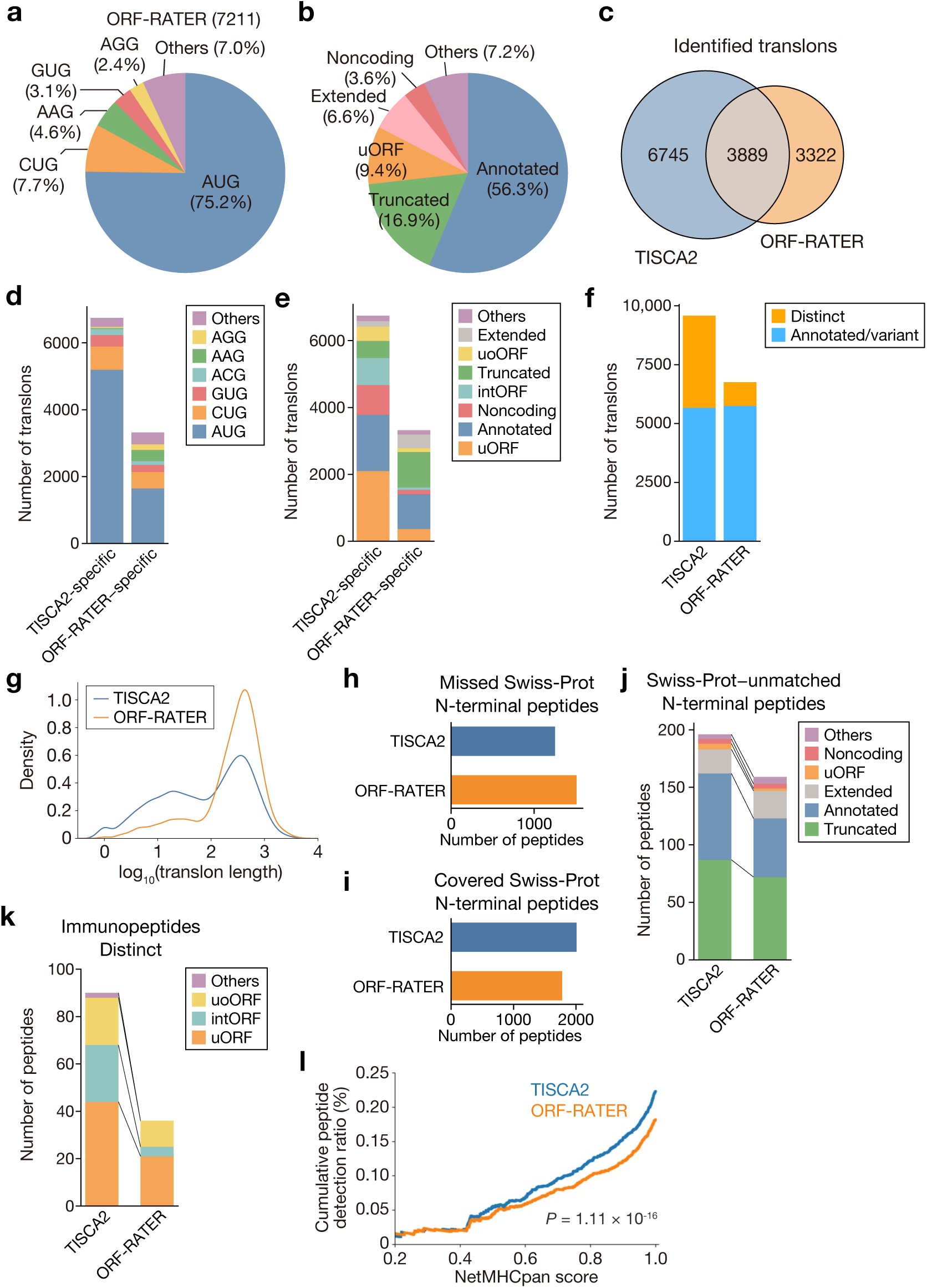
TISCA2 improves detection of distinct translons. **a**, Start-codon composition of 7,211 translons identified by ORF-RATER in HEK293T cells. **b**, Classification of ORF-RATER-predicted translons by types. **c**, Overlap between TISCA2- and ORF-RATER-defined translons. **d**, Start-codon composition of TISCA2-specific and ORF-RATER-specific translons. **e**, Classification of TISCA2-specific and ORF-RATER-specific translons by types. **f**, Numbers of annotated/variant and distinct translons identified by TISCA2 and ORF-RATER. **g**, Length distributions of translons identified by TISCA2 and ORF-RATER. **h**, Numbers of Swiss-Prot–derived N-terminal peptides not represented by the TISCA2 or ORF-RATER prediction database (missed Swiss-Prot N-terminal peptides). **i**, Numbers of Swiss-Prot–derived N-terminal peptides represented by each prediction database (covered Swiss-Prot N-terminal peptides). **j**, Swiss-Prot–unmatched N-terminal peptides identified using the TISCA2- or ORF-RATER-derived databases, classified according to translon types. **k**, HLA class I immunopeptides derived from distinct translons identified using the TISCA2- or ORF-RATER-derived databases, classified according to translon types. **l**, Cumulative detection ratio of HLA class I immunopeptides as a function of NetMHCpan raw score for TISCA2- and ORF-RATER-defined translons. *P* value was calculated using Hotelling’s *T*-squared test.

Translon classification showed that ORF-RATER identified higher proportions of annotated CDSs and variant translons, including truncated and extended CDSs, than TISCA2 (Fig. 2b).

The two approaches overlapped only partially. TISCA2 identified 10,634 TISs, whereas ORF-RATER identified 7,211 translons, with 3,889 start sites shared between the two methods (Fig. 2c). The common set was dominated by AUG-initiated annotated CDSs and uORFs (Extended Data Fig. 3b), whereas method-specific translons showed distinct features. TISCA2-specific translons were enriched for AUG initiation, whereas ORF-RATER-specific translons showed a higher fraction of near-cognate start codons (Fig. 2d). TISCA2-specific translons occurred in 5′ UTRs (uORFs), within CDSs (intORFs and truncated CDSs), and in noncoding transcripts, whereas ORF-RATER-specific translons were particularly enriched for truncated CDSs (Fig. 2e). Comparison of these classes showed that TISCA2 identified more distinct translons, including uORFs, uoORFs and intORFs, whereas ORF-RATER preferentially identified annotated and variant translons (Fig. 2f). Consistent with this difference, TISCA2-defined translons were shorter than ORF-RATER-defined translons (Fig. 2g). These trends were not explained simply by cutoff selection, as increasing the cosine-similarity cutoff for TISCA2 or the ORF-RATER score cutoff progressively reduced the number of calls and increased the fraction of annotated CDSs without reversing the overall class distributions (Extended Data Fig. 3c).

To orthogonally evaluate these predictions, we performed HUNTER N-terminomics and searched the data against combined databases containing Swiss-Prot proteins together with products of translons predicted by either TISCA2 or ORF-RATER^48^. Swiss-Prot-derived N-terminal peptides were classified as missed or covered according to whether they were absent from or represented in the corresponding prediction set, respectively^49^. TISCA2 missed fewer Swiss-Prot N-terminal peptides than ORF-RATER (Fig. 2h) and covered more Swiss-Prot N-terminal peptides (Fig. 2i), indicating broader recovery of curated protein N-termini. We next focused on Swiss-Prot–unmatched N-terminal peptides, defined as peptides that were absent from Swiss-Prot but identified from the TISCA2 or ORF-RATER prediction databases. This class includes GENCODE-annotated products that are not curated in Swiss-Prot, as well as products of translons not annotated as CDSs. TISCA2 yielded more Swiss-Prot–unmatched N-terminal peptide identifications than ORF-RATER (Fig. 2j). Notably, although TISCA2 predicted fewer truncated translons than ORF-RATER (Figs. 1f and 2b), it recovered more N-terminal peptides assigned to truncated products (Fig. 2j). At the *HNRNPU* locus, TISCA2 identified an intORF supported by an N-terminal peptide, whereas this region lacked a prominent TI-seq peak and was not detected by ORF-RATER (Extended Data Fig. 3d).

Most N-terminal peptides assigned to TISCA2- or ORF-RATER-predicted products were derived from truncated or extended CDSs, whereas peptides supporting other translons (e.g. uORFs and intORFs) were much rarer (Fig. 2j). This is consistent with the difficulty of detecting small translon-derived products by conventional mass spectrometry, because short translons generate fewer uniquely assignable peptides^17,50^.

Total proteome analysis similarly detected only a limited number of peptides derived from distinct translons, although more distinct translons received proteomic support using TISCA2 predictions than ORF-RATER predictions (Extended Data Fig. 3e).

We therefore asked whether HLA class I immunopeptidomics could provide complementary evidence for products of distinct translons. Because HLA-I peptidomics directly profiles naturally presented peptides and does not require tryptic digestion, it can capture peptides derived from short translons^51–53^. Proteogenomic searches of HLA-I immunopeptidomics data identified more HLA-I peptides derived from distinct translons using TISCA2 predictions than ORF-RATER predictions (Fig. 2k). The length distribution of TISCA2-derived HLA-I peptides was consistent with typical MHC-I-presented peptides, with most peptides being 8–11 amino acids long (Extended Data Fig. 3f).

Finally, we evaluated the antigen-presentation potential of peptides derived from predicted translons using NetMHCpan raw scores^54^. Across the intermediate-to-high NetMHCpan score range, peptides derived from TISCA2-predicted translons yielded a higher cumulative peptide detection ratio than peptides derived from ORF-RATER-predicted translons (Fig. 2l). Thus, TISCA2 improves the detection of distinct translons and provides broader coverage of experimentally detected HLA-I-presented peptides. Together, these results indicate that TISCA2 complements and extends Ribo-seq/TI-seq–based approaches by improving the detection of short translons overlapping CDSs.

### Arsenite increases downstream-initiated translation

To determine whether the ISR broadly remodels start-site selection, we performed eIF3D and eIF5B Sel-TCP-seq in arsenite-treated HEK293T cells. We refer to all translons initiated downstream of the annotated start codon as downstream-initiated translons. Metagene analysis showed that arsenite treatment extended the distribution of eIF3D footprints from annotated 5′ UTRs into CDSs and annotated 3′ UTRs. eIF5B footprints retained sharp enrichment around annotated start codons, but arsenite treatment also increased eIF5B-associated signals within CDSs and annotated 3′ UTRs (Fig. 3a). These patterns suggested that stress increases scanning beyond upstream start sites and promotes downstream initiation. To quantify this effect, we calculated a leaky-scanning score from eIF3D and eIF5B footprints, defined as the relative abundance of CDS-proximal footprints compared with start-codon–proximal footprints^55^. Arsenite treatment significantly increased the leaky-scanning score for both eIF3D and eIF5B footprints (Fig. 3b), suggesting a global shift in start-site selection during the ISR.

**Fig. 3:**
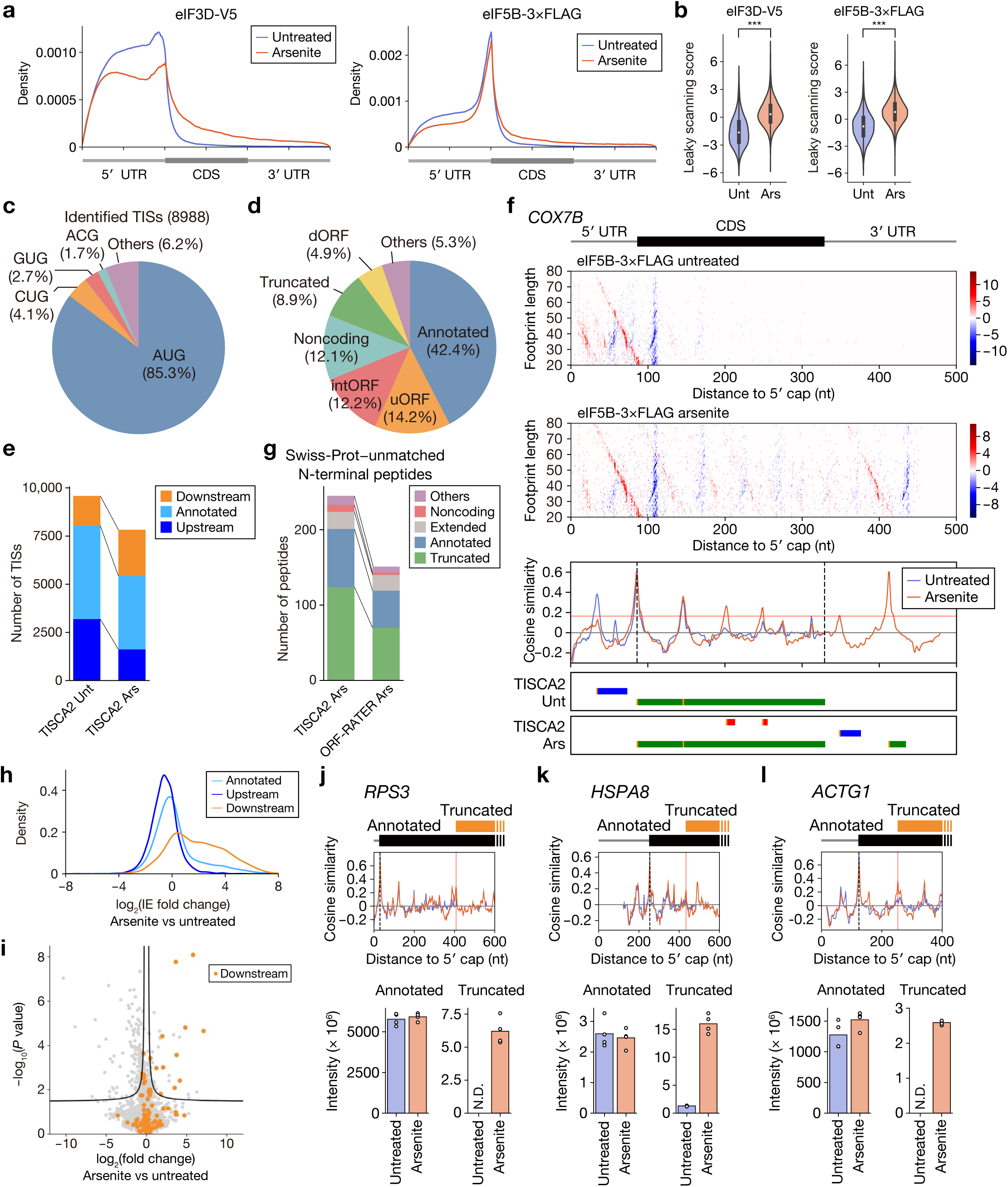
Stress shifts translation initiation downstream. **a**, Metagene profiles of eIF3D-V5 and eIF5B-3×FLAG Sel-TCP-seq footprints in untreated and arsenite-treated HEK293T cells. Representative data of two biological replicates are shown. All reads were mapped to all human protein-coding transcripts with a 5′-UTR and 3′-UTR of >100 nt (*n* = 52,692). **b**, Leaky-scanning scores calculated from eIF3D-V5 and eIF5B-3×FLAG footprints in untreated and arsenite-treated cells. \*\*\**P* < 0.001, Mann–Whitney *U* test. **c**, Start-codon composition of 8,988 TISs identified by TISCA2 after arsenite treatment. **d**, Classification of translons identified after arsenite treatment by types. **e**, Numbers of upstream, canonical and downstream TISs identified by TISCA2 in untreated and arsenite-treated cells. **f**, Representative TISCA2 analysis of the *COX7B* transcript in untreated and arsenite-treated cells. eIF5B-3×FLAG differential footprint plots and cosine-similarity profiles are shown. Black dotted lines indicate the canonical start codon and stop codon. Horizontal red line indicates the cutoff value (0.164). Horizontal bars indicate the three translation reading frames, and thin vertical yellow regions indicate TISs identified by TISCA2. **g**, Swiss-Prot-unmatched N-terminal peptides identified using TISCA2- or ORF-RATER-derived databases after arsenite treatment, classified according to translon category types. **h**, Distributions of IE fold changes following arsenite treatment for canonical, upstream and downstream TISs. **i**, Differential N-terminomics analysis of all identified N-terminal peptides in untreated and arsenite-treated cells (*n* = 4 biologically independent samples). Fold changes in peptide abundance are plotted against *P* values, and peptides derived from downstream-initiated translon products are highlighted. Black curves indicate 1% FDR (false discovery rate). *P* value was calculated using Student’s *t* test. **j–l**, TISCA2 profiles and quantitative N-terminal proteomics of annotated canonical and truncated products from *RPS3* (**j**), *HSPA8* (**k**) and *ACTG1* (**l**) in untreated and arsenite-treated cells. Individual biological replicates are shown as points (*n* = 4 biologically independent samples). N.D., not detected.

We next applied TISCA2 to arsenite-treated cells. The characteristic eIF5B metagene and differential-metagene patterns were retained under arsenite treatment (Extended Data Fig. 4a), and a condition-specific cosine-similarity cutoff was determined (Extended Data Fig. 4b). TISCA2 identified 8,988 TISs after arsenite treatment. The start-codon composition was broadly similar to that observed in untreated cells, with AUG accounting for 85.3% of TISs, followed by CUG, GUG and ACG (Fig. 3c). In contrast, the distribution of start sites along transcripts changed markedly. Arsenite treatment reduced the proportion of uORFs and increased the proportions of intORFs, truncated translon variants and dORFs, indicating a shift toward downstream initiation (Fig. 3d). Comparison of TISs identified in untreated and arsenite-treated cells further showed that many TISs were detected in only one condition (Extended Data Fig. 4c). When TISs were grouped by their position relative to the annotated start codon, arsenite treatment reduced upstream TISs and increased downstream TISs (Fig. 3e).

This redistribution of initiation was also evident at individual loci. At *COX7B*, TISCA2 detected the annotated CDS TIS and an additional upstream TIS under untreated conditions, whereas arsenite treatment induced additional TISCA2 peaks within the CDS and annotated 3′ UTR (Fig. 3f). Ribo-seq signals supported translation of the corresponding downstream regions, including translons mapping to the annotated 3′ UTR (Extended Data Fig. 4d). Conversely, the upstream TIS detected in *COX7B* under untreated conditions no longer met the TISCA2 calling criteria after arsenite treatment. Comparison with ORF-RATER under arsenite treatment reproduced the major differences observed in untreated cells: TISCA2 identified a larger fraction of distinct translons, whereas ORF-RATER favored annotated and variant translons (Extended Data Fig. 4e–i).

We then performed HUNTER N-terminomics to assess translon products induced or detected under stress. Under arsenite treatment, searches using the TISCA2-derived database identified more Swiss-Prot–unmatched N-terminal peptides than searches using the ORF-RATER-derived database (Fig. 3g). Notably, TISCA2 recovered more N-terminal peptides assigned to truncated CDSs, supporting the production of a subset of downstream-initiated translon products under stress.

We next calculated IE for TISs identified in either untreated or arsenite-treated cells and compared IE between conditions. Arsenite treatment globally reduced IE at annotated CDS starts and upstream TISs, whereas IE increased at downstream TISs (Fig. 3h). At *ATF4*, a well-established ISR target, TISCA2 detected increased IE at the annotated ATF4 start site after arsenite treatment, confirming that the method captures known stress-dependent translational regulation (Extended Data Fig. 4j). Finally, we used N-terminomics to examine quantitative changes in downstream-initiated translon products after arsenite treatment. Although most downstream-initiated N-terminal peptides did not show statistically significant changes, most significantly altered peptides increased in abundance after arsenite treatment (Fig. 3i). At individual loci, N-terminal peptides corresponding to truncated CDSs of *RPS3*, *HSPA8* and *ACTG1* showed marked increases in abundance after arsenite treatment (Fig. 3j–l). In contrast, N-terminal peptides corresponding to the annotated CDS start sites of the same genes did not show substantial increases (Fig. 3j–l).

Together, these findings show that the arsenite-induced ISR broadly remodels start-site selection across the transcriptome. These data extend the classical uORF-centered view of stress-regulated translation and reveal that the ISR promotes translation initiation downstream of annotated CDS start sites.

### Sequence, structural and proteomic features of TISCA2-predicted translon products

TISCA2 improved the detection of distinct translons overlapping annotated CDSs, including uoORFs and intORFs. We therefore investigated the sequence and structural properties of products predicted from TISCA2-specific translons. For each TISCA2-specific translon-derived product, we divided the predicted amino acid sequence into overlapping 20-amino-acid windows, shifted by one residue, and determined whether each window exactly matched any sequence in the human Swiss-Prot database. We defined the 20-amino-acid exact-match fraction as the proportion of windows with a 100% sequence match to a Swiss-Prot protein. For subsequent analyses, we selected predicted products with a 20-amino-acid exact-match fraction of 0.1 or less (Fig. 4a). Products of 20 amino acids or fewer were excluded from subsequent analyses.

**Fig. 4:**
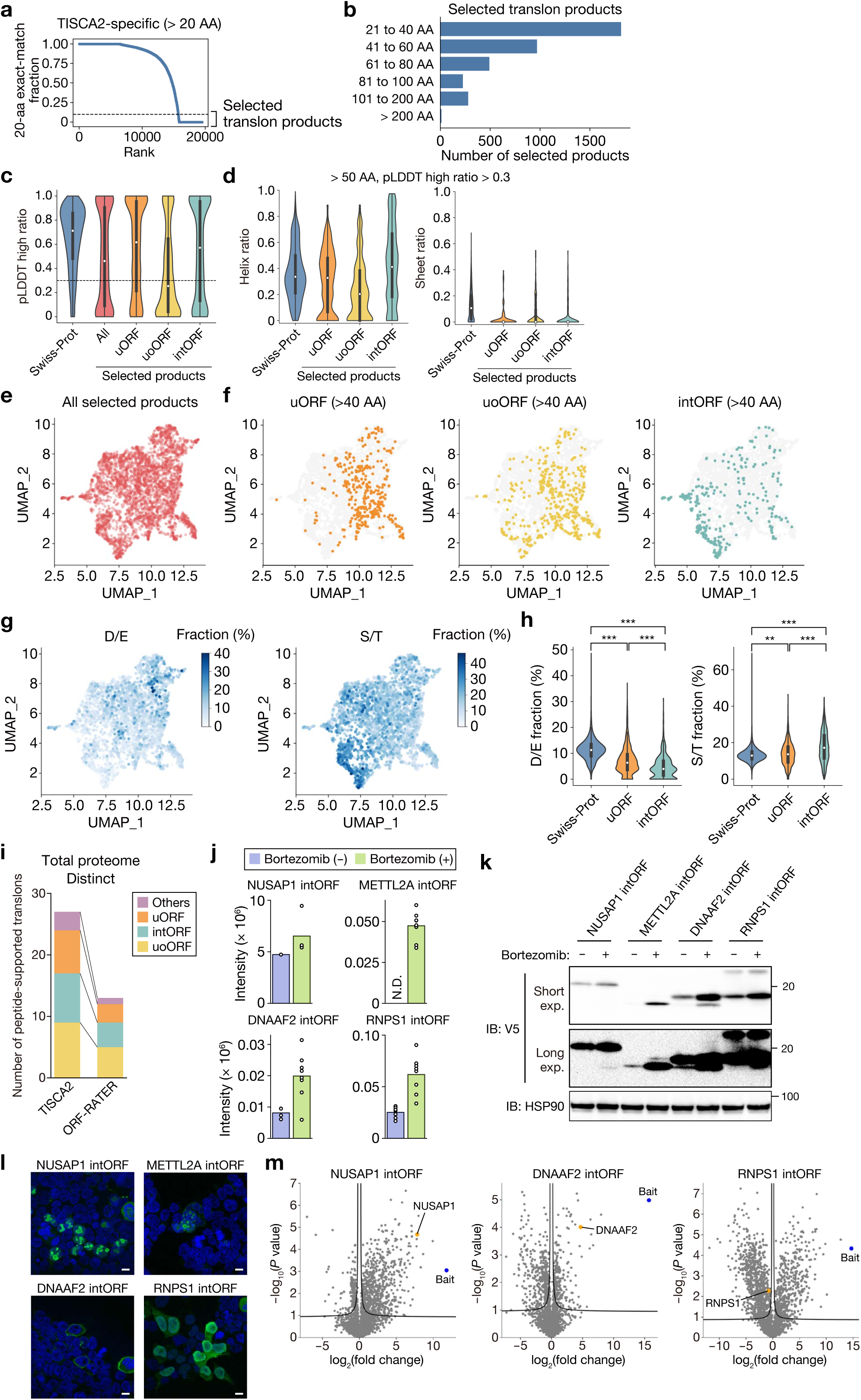
Sequence, structural and cellular characterization of TISCA2-defined translon products. **a**, TISCA2-specific translon-derived products longer than 20 amino acids ranked by their 20-amino-acid exact-match fraction against human Swiss-Prot proteins. Products with an exact-match fraction ≤0.1 were classified as selected translon products. **b**, Length distribution of selected translon products. **c**, Distribution of pLDDT-high ratios for Swiss-Prot proteins and the indicated products classified by translon types. The pLDDT-high ratio was defined as the fraction of residues with a pLDDT score >70. **d**, Distribution of ratios of helices or sheets for Swiss-Prot proteins and the indicated products classified by translon types. **e**, UMAP visualization of ESM-2 sequence representations of all selected products. **f**, Distribution of uORF-, uoORF- and intORF-derived products longer than 40 amino acids in the UMAP space shown in **e**. **g**, UMAP plots colored according to the fraction of aspartate/glutamate (D/E) or serine/threonine (S/T) residues. **h**, Distributions of D/E and S/T fractions in Swiss-Prot proteins and uORF- and intORF-derived products. \*\**P* < 0.01, \*\*\**P* < 0.001, Steel-Dwass test. **i**, Distinct translons supported by total proteomics using TISCA2- or ORF-RATER-derived search databases, classified according to translon types. **j**, Quantification of unique peptides supporting the indicated intORF-derived products in untreated and Bortezomib (Btz)-treated HEK293T cells. **k**, Immunoblot analysis of V5-tagged NUSAP1, METTL2A, DNAAF2 and RNPS1 intORF-derived products expressed in HEK293T cells with or without Btz treatment. HSP90 was used as a loading control. A longer exposure of the V5 immunoblot is shown. **l**, Representative immunofluorescence images of the indicated V5-tagged intORF-derived products in HEK293T cells. Nuclei were counterstained with DAPI (blue) and V5-tagged proteins were detected with an anti-V5 antibody (green). Scale bars, 10 μm. **m**, Proteins associated with the indicated intORF-derived products identified by V5 immunoprecipitation followed by mass spectrometry (*n* = 3 biologically independent samples). N.D., not detected.

Most selected products were short, with the largest group comprising products of 21–40 amino acids. Products longer than 100 amino acids were also identified, although those exceeding 200 amino acids were uncommon (Fig. 4b). We next used AlphaFold 3 (AF3) to predict their structures^56^. To summarize local prediction confidence, we defined the pLDDT-high ratio as the fraction of residues with a pLDDT score greater than 70. Although many of these products yielded low-confidence AF3 predictions, a subset showed high local confidence (Fig. 4c). High pLDDT-high ratios became less frequent when the analysis was restricted to products longer than 50 amino acids (Extended Data Fig. 5a). To characterize secondary structure, we applied Dictionary of Protein Secondary Structure (DSSP) to structural models of selected products longer than 50 amino acids with a pLDDT-high ratio greater than 0.3 and calculated the fractions of residues assigned to helices and β-sheets. intORF-derived products showed higher helix ratios than uORF- and uoORF-derived products and Swiss-Prot proteins, whereas β-sheet ratios were generally low across all groups, including Swiss-Prot proteins (Fig. 4d).

To further characterize these products, we extracted sequence representations using the protein language model Evolutionary Scale Modeling 2 (ESM-2) and visualized them using Uniform Manifold Approximation and Projection (UMAP) (Fig. 4e)^57^. Among products longer than 40 amino acids, uORF- and intORF-derived products occupied distinct regions of the ESM-2 sequence space, whereas uoORF-derived products were more broadly distributed (Fig. 4f). Similar class-dependent differences were also observed among products of 21–40 amino acids, although the separation was more pronounced for products longer than 40 amino acids (Extended Data Fig. 5b).

We next investigated sequence features associated with this separation. Mapping amino-acid composition onto the UMAP revealed distinct distributions of acidic and serine/threonine-rich sequences (Fig. 4g). uORF-derived products contained a higher fraction of aspartate and glutamate residues (D/E) than intORF-derived products, although their D/E content remained lower than that of Swiss-Prot proteins. In contrast, intORF-derived products showed a higher fraction of serine and threonine residues (S/T) than either uORF-derived products or Swiss-Prot proteins (Fig. 4h). These compositional differences may partly reflect sequence constraints associated with overlap with annotated CDSs and the translated reading frames.

Products of unannotated translons are often unstable and can be targeted by proteasome-dependent quality-control pathways^58^. To assess the proteomic support and proteasome sensitivity of TISCA2-defined products, we treated HEK293T cells with the proteasome inhibitor bortezomib (Btz), acquired total proteome data using an Orbitrap Astral mass spectrometer and performed proteogenomic searches against databases containing Swiss-Prot proteins together with products predicted by either TISCA2 or ORF-RATER (Extended Data Fig. 5c). Searches using the TISCA2-derived database provided proteomic support for more distinct translons, including intORFs, than searches using the ORF-RATER-derived database (Fig. 4i and Extended Data Fig. 5d,e). Most detected products were supported in both untreated and Btz-treated cells, whereas a subset was detected preferentially or exclusively after Btz treatment, suggesting that some TISCA2-defined products are proteasome-sensitive.

For selected products detected by total proteomics, we next compared Btz-dependent changes in abundance with the AF3-derived pLDDT-high ratio (Extended Data Fig. 5f). Btz responsiveness showed a partial correspondence with the pLDDT-high ratio: several products with low-confidence structural predictions accumulated after proteasome inhibition, although this relationship was not consistent across all products (Extended Data Fig. 5f). Thus, predicted structural confidence alone was insufficient to explain proteasome sensitivity.

### Selected intORF-derived products exhibit distinct cellular properties

Because TISCA2 identified numerous intORFs, including intORFs with increased initiation under stress and amino-acid compositions distinct from those of uORF-derived products, we next examined selected intORF-derived products individually. We focused on intORFs in *NUSAP1*, *METTL2A*, *DNAAF2* and *RNPS1*, encoding predicted products of 156, 77, 112 and 136 amino acids, respectively, each of which was supported by unique peptides in the total proteome data (Fig. 4j). The abundances of the supporting peptides showed varying responses to Btz, with peptides from the *METTL2A* intORF-derived product showing the strongest enrichment. We then tested whether this behavior was recapitulated by V5-tagged products in HEK293T cells. Consistent with the total proteomics data, all four V5-tagged products increased to varying degrees following Btz treatment (Fig. 4k). The *METTL2A* intORF-derived product was particularly sensitive to proteasome inhibition, showing only weak accumulation in untreated cells but a marked increase after Btz treatment. AF3 models showed generally low-to-moderate pLDDT scores, with localized regions of higher confidence in some products (Extended Data Fig. 5g).

Immunofluorescence analysis revealed strikingly distinct subcellular localization patterns among the four products (Fig. 4l). The *NUSAP1* and *METTL2A* intORF-derived products were enriched in nucleolar regions, whereas the *DNAAF2* intORF-derived product showed prominent localization at the cell periphery, consistent with plasma-membrane localization. In contrast, the *RNPS1* intORF-derived product displayed a broader distribution across the nucleus and cytoplasm.

We further examined proteins associated with the intORF-derived products by V5 immunoprecipitation followed by mass spectrometry. The *METTL2A* intORF-derived product was excluded from mass spectrometric analysis because of its low expression. The *NUSAP1* and *DNAAF2* intORF-derived products co-purified with the corresponding annotated CDS-derived proteins, whereas enrichment of the annotated RNPS1 protein was not detected with the *RNPS1* intORF-derived product (Fig. 4m).

Gene Ontology enrichment analysis of the co-purifying proteins revealed rRNA-processing- and ribosome-biogenesis-related categories for the *NUSAP1* intORF-derived product, consistent with its enrichment in nucleolar regions (Fig. 4l and Extended Data Fig. 5h). Protein-folding-related categories were enriched for the *DNAAF2* intORF-derived product, whereas translation- and transcriptional-regulation-related categories were enriched for the *RNPS1* intORF-derived product (Extended Data Fig. 5h). This enrichment pattern was consistent with the distribution of the *RNPS1* intORF-derived product across the nucleus and cytoplasm. Together, these results identify protein associations that provide clues to the potential cellular roles of these intORF-derived products.

### TISCA2 reveals tissue-specific translation initiation landscapes in vivo

Because inhibitor-based TIS profiling in barrier-protected tissues can be complicated by delayed drug penetration^31^, we next asked whether TISCA2 could be applied directly to mammalian tissues. We generated eIF5B-3×FLAG knock-in mice and performed eIF5B Sel-TCP-seq in perfusion-fixed brain, kidney and testis (Extended Data Fig. 6a). In all three tissues, eIF5B-associated footprints showed clear enrichment around annotated start codons, recapitulating the TIS-associated pattern observed in cultured cells (Fig. 5a and Extended Data Fig. 6b).

**Fig. 5:**
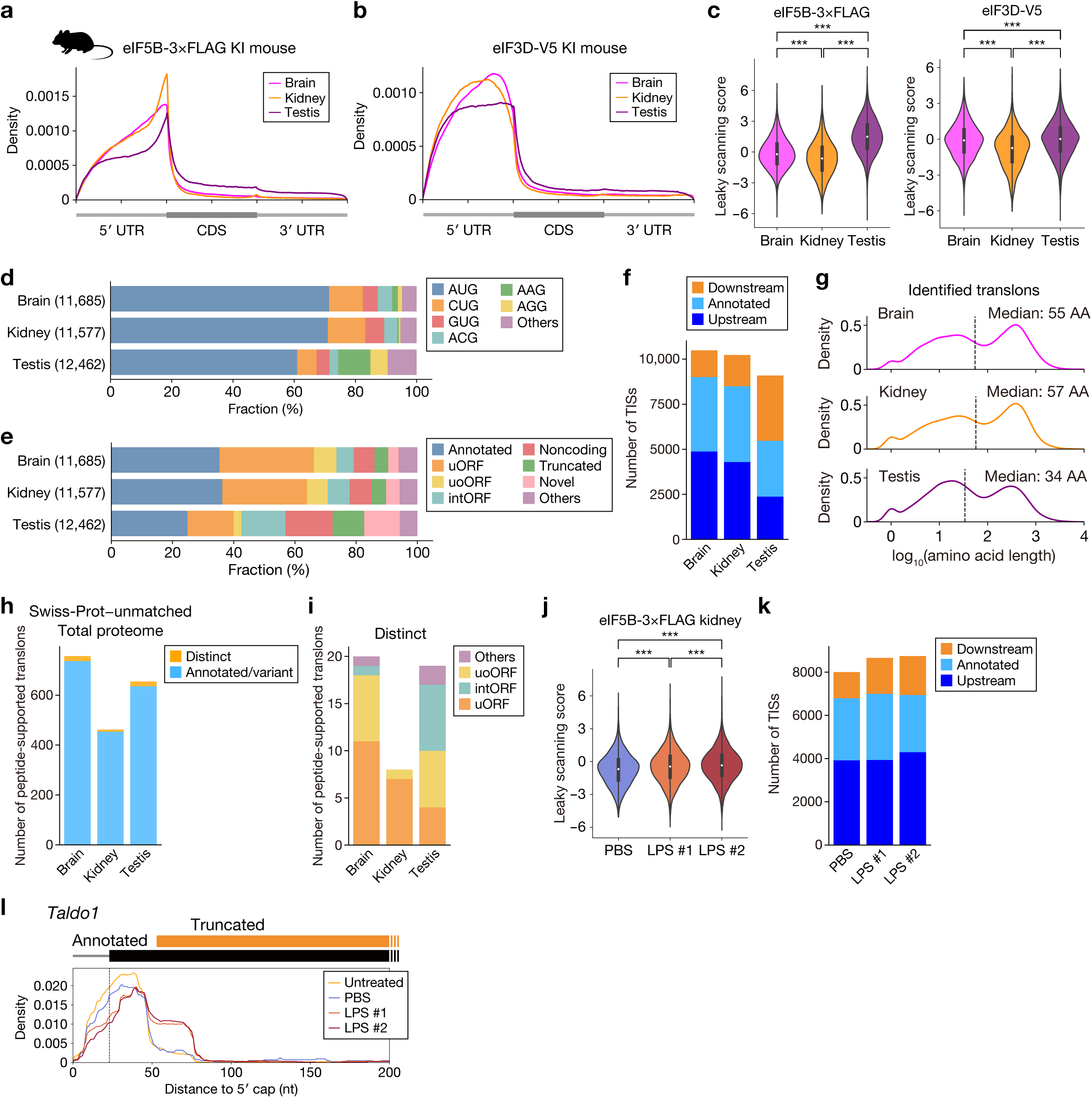
TISCA2 reveals tissue-specific translation initiation landscapes in vivo. **a**,**b**, Metagene profiles of eIF5B-3×FLAG (**a**) or eIF3D-V5 (**b**) Sel-TCP-seq footprints in mouse brain, kidney and testis. Representative data of two biological replicates are shown for eIF5B-3×FLAG in brain and eIF3D-V5. All reads were mapped to all mouse protein-coding transcripts with a 5′-UTR and 3′-UTR of >100 nt (*n* = 31,343). **c**, Leaky-scanning scores calculated from eIF5B-3×FLAG and eIF3D-V5 footprints in the indicated tissues. \*\*\**P* < 0.001, Steel-Dwass test. **d**, Start-codon composition of TISs identified by TISCA2 in brain, kidney and testis. The number of identified TISs is indicated. **e**, Classification of translons identified in the indicated tissues by types. **f**, Numbers of upstream, canonical and downstream TISs identified in brain, kidney and testis. **g**, Length distributions of translon products predicted by TISCA2 in the indicated tissues. Median lengths are indicated. **h**, Numbers of Swiss-Prot–unmatched annotated/variant and distinct translons supported by total proteomics in brain, kidney and testis. **i**, Distinct translons supported by total proteomics, classified according to translon types. **j**, Leaky-scanning scores calculated from eIF5B-3×FLAG footprints in kidney collected from mice under indicated conditions. \*\*\**P* < 0.001, Steel-Dwass test. **k**, Numbers of upstream, canonical and downstream TISs identified in kidney collected from mice under indicated conditions. **l**, Read aggregation plots of eIF5B footprints in kidney collected from mice under indicated conditions for *Taldo1*.

Notably, eIF5B-associated footprints in the testis extended further downstream of annotated start codons than those in brain or kidney (Fig. 5a). To independently examine scanning ribosomes, we generated eIF3D-V5 knock-in mice and performed eIF3D Sel-TCP-seq in the same tissues. eIF3D footprints similarly extended further downstream in the testis (Fig. 5b). Consistent with these patterns, the leaky-scanning score was the highest in the testis for both eIF3D- and eIF5B-associated footprints (Fig. 5c). These concordant profiles are consistent with increased ribosomal scanning beyond annotated start sites.

We next applied TISCA2 to the tissue eIF5B Sel-TCP-seq data. TISCA2 identified 11,685 TISs in brain, 11,577 in kidney and 12,462 in testis (Fig. 5d,e and Extended Data Fig. 6c). Brain and kidney showed start-codon distributions broadly similar to that observed in HEK293T cells, with AUG predominating and CUG, GUG and ACG accounting for most near-cognate initiation events (Fig. 5d). The population of unannotated translons was also dominated by uORFs (Fig. 5e). By contrast, the testis showed a distinct initiation landscape. The fraction of AUG-initiated TISs was reduced, whereas candidate TISs at AAG and AGG codons were markedly increased (Fig. 5d).

The testis also contained higher proportions of intORFs and truncated ORFs than brain or kidney (Fig. 5e). Notably, AAG- and AGG-associated candidate TISs in the testis were predominantly located downstream of annotated start codons (Extended Data Fig. 6d), although whether these signals represent genuine initiation remains uncertain.

Consistent with these differences, positional classification of TISs revealed a higher proportion of downstream candidate TISs in the testis (Fig. 5f). Predicted translon products were also shorter, with median lengths of 55, 57 and 34 amino acids in brain, kidney and testis, respectively (Fig. 5g). When the analysis was restricted to TISCA2-predicted truncated ORFs, their lengths relative to the corresponding annotated products also showed a broader shift toward shorter products in the testis (Extended Data Fig. 6e).

We next sought orthogonal proteomic evidence for this tissue-specific pattern. Proteogenomic analysis of total proteomes identified Swiss-Prot-unmatched products in all three tissues (Fig. 5h). Among products of distinct translons, the testis showed a greater contribution from intORF-derived products than brain or kidney (Fig. 5i). Because conventional bottom-up proteomics cannot readily distinguish truncated products from their corresponding annotated proteins when they share internal peptides, we further analyzed N-terminal peptides. Swiss-Prot-unmatched N-terminal peptides mapped to predicted products that showed a shift toward shorter lengths in the testis (Extended Data Fig. 6f,g). A similar shift was observed when the analysis was restricted to truncated CDSs (Extended Data Fig. 6h). Moreover, when the lengths of N-terminally detected truncated products were normalized to those of their corresponding annotated products, the testis again showed a broader distribution toward shorter relative lengths (Extended Data Fig. 6i). Thus, the N-terminal proteomics results were consistent with the shift toward shorter products predicted by TISCA2 in the testis.

### Inflammatory and chronic restraint stress alter start-site selection in vivo

To examine whether the redistribution of initiation observed after arsenite treatment also occurs in an in vivo inflammatory setting, we used an LPS-induced endotoxemia model previously shown to induce renal eIF2α phosphorylation and translational repression^59,60^. LPS treatment increased renal eIF2α phosphorylation and the expression of the inflammatory genes *Il1b* and *Il6*, supporting activation of inflammatory and ISR-associated responses under these conditions (Extended Data Fig. 7a,b). TISCA2 analysis revealed increased leaky-scanning scores and increased numbers of downstream TISs in kidneys from two independently treated mice compared with the PBS control (Fig. 5j,k). At *Taldo1*, LPS treatment increased eIF5B-associated signals near internal start sites corresponding to predicted N-terminally truncated products (Fig. 5l). These results suggest that inflammatory stress can also alter start-site selection in vivo, with increased use of downstream initiation sites.

We next asked whether start-site selection was also altered in a distinct model of chronic behavioral stress. Mice were subjected to chronic restraint stress (CRS), after which brains were analyzed by TISCA2 (Extended Data Fig. 7c). CRS increased the leaky-scanning score calculated from eIF5B-associated footprints (Extended Data Fig. 7d). However, the numbers of upstream, annotated and downstream TISs were broadly similar between control and CRS-treated brains, with no consistent increase in downstream TISs (Extended Data Fig. 7e).

We therefore examined changes in initiation efficiency at individual TISs. Representative transcripts, including *Hnrnpu* and *Ids*, showed differential changes among TISs within the same transcript, with reduced initiation at upstream near-cognate sites relative to downstream AUG sites after CRS (Extended Data Fig. 7f). Genome-wide, AUG-initiated TISs shifted toward increased IE, whereas near-cognate TISs shifted toward reduced IE (Extended Data Fig. 7g). Positional classification showed a corresponding difference: annotated TISs tended toward increased IE, whereas upstream TISs tended toward reduced IE following CRS (Extended Data Fig. 7h). These concordant changes are consistent with increased stringency of start-codon selection under chronic stress, favoring initiation at AUG codons over alternative upstream and near-cognate sites^61^.

Genes showing increased IE at annotated TISs were enriched for functions related to cytoplasmic translation, pre- and postsynaptic translation, mitochondrial ATP synthesis, aerobic respiration and protein transport (Extended Data Fig. 7i), suggesting coordinated translational adaptation to chronic stress. Thus, whereas LPS treatment increased the number of downstream TISs in the kidney, CRS primarily altered the relative efficiency of existing TISs in the brain, particularly reducing initiation at upstream and near-cognate sites.

### Development of IncepTIS for condition-dependent TIS prediction

TISCA2 profiles from untreated and arsenite-treated HEK293T cells provide experimental training targets for condition-dependent initiation prediction. We therefore developed IncepTIS, a deep learning model that predicts position-resolved TISCA2 cosine-similarity profiles from transcript sequence and experimental condition. Using measured profiles as training targets allows the model to learn initiation signals beyond annotated start codons. IncepTIS combines sequence representations extracted by Nucleotide Transformer v3 (NTv3) with a condition indicator and uses an Attention U-Net prediction head to generate nucleotide-resolution predictions^62,63^ (Fig. 6a and Extended Data Fig. 8a). The condition indicator enables the model to generate separate predictions for the same transcript under untreated and arsenite-stress conditions.

**Fig. 6:**
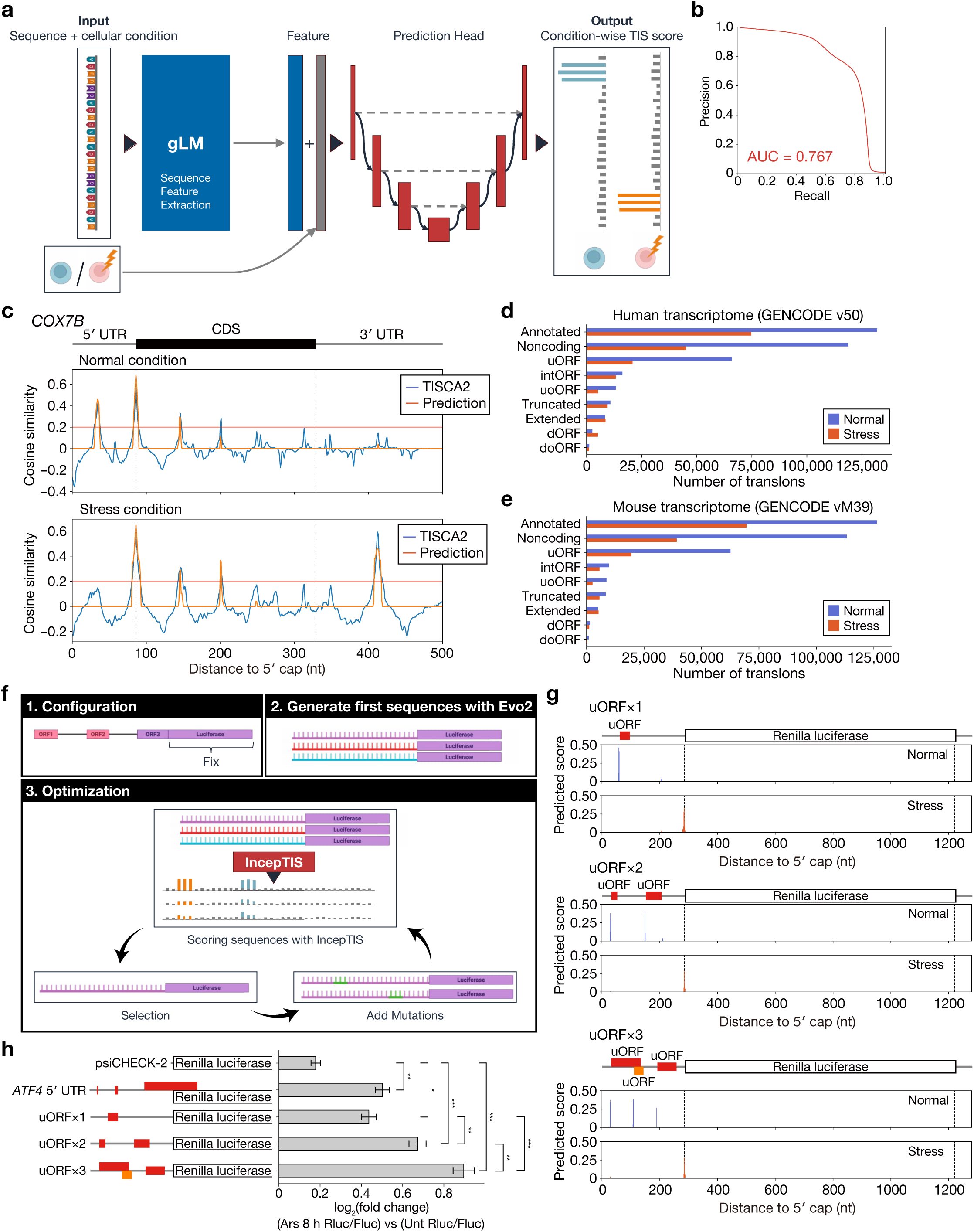
IncepTIS predicts condition-dependent translation initiation landscapes and guides sequence design. **a,** Overview of IncepTIS. Transcript sequences are encoded by Nucleotide Transformer v3 (NTv3), combined with cellular state and passed to an Attention U-Net to predict condition-specific TISCA2 cosine similarity profiles. **b,** Precision–recall curve pooling held-out predictions from transcript-level 10-fold cross-validation. Positions with measured TISCA2 cosine similarity ≥0.2 were defined as positive. PR-AUC, 0.767. **c,** Measured TISCA2 profiles and IncepTIS predictions for the held-out *COX7B* transcript in untreated (top) and arsenite-treated (bottom) HEK293T cells. Vertical dashed lines mark annotated CDS boundaries; horizontal red lines indicate a cosine similarity of 0.2. **d,e,** Numbers of predicted translons by annotation class across GENCODE human (Release 50; **d**) and mouse (Release M39; **e**) transcripts under untreated and arsenite-stress conditions. Predictions in mouse were not experimentally validated. **f,** Overview of IncepTIS-evolve. Initial sequences generated using Evo 2 are optimized by a genetic algorithm using condition-specific IncepTIS predictions. The 5′ leader is mutable, whereas the Renilla luciferase start codon and coding sequence are fixed. Optimization favors initiation at specified upstream sites under untreated conditions and at the luciferase start site under stress. **g,** IncepTIS-predicted initiation profiles under untreated and arsenite-stress conditions for three independently designed 5′ leaders targeting one, two or three uORF initiation sites (uORFx1, uORFx2 and uORFx3). Predicted uORFs and the Renilla luciferase CDS are indicated. **h,** Dual-luciferase reporter assays of the three designed leaders, the ATF4 5′ UTR and the parental psiCHECK-2 vector in HEK293T cells with or without sodium arsenite treatment (40 μM, 8 h). Values represent log_2_ fold changes in the Renilla-to-firefly luciferase activity ratio after arsenite treatment relative to untreated cells. Data are mean ± s.d.; n = 3 independent biological replicates. Statistical significance was assessed by one-way analysis of variance followed by Tukey’s multiple-comparisons test. \**P* < 0.05, \*\**P* < 0.01, \*\*\**P* < 0.001.

Training jointly optimized agreement with measured cosine-similarity values and classification of high-scoring positions (see Methods). Training and validation loss trajectories are shown in Extended Data Fig. 8b,c. Validation PR-AUC increased during training and subsequently plateaued, with similar trends across folds (Extended Data Fig. 8d,e). In transcript-level 10-fold cross-validation, IncepTIS achieved a pooled nucleotide-level PR-AUC of 0.767 against TISCA2-derived labels (mean ± s.d. across folds, 0.767 ± 0.004), with consistent performance across folds (Fig. 6b and Extended Data Fig. 8f).

IncepTIS also captured condition-dependent features of initiation profiles. At COX7B, a transcript held out from training, the model reproduced the reduction in upstream initiation and the appearance of downstream initiation peaks under arsenite treatment, including a peak within the annotated 3′ UTR (Fig. 6c). These predictions were consistent with the experimentally measured TISCA2 profiles, illustrating the ability of IncepTIS to predict condition-dependent changes in start-site usage.

We next applied IncepTIS to GENCODE human transcripts and extended prediction to mouse transcripts to generate an atlas of predicted initiation under untreated and arsenite-stress conditions (Fig. 6d,e). Predictions encompassed annotated CDSs and diverse additional translon classes, including uORFs, intORFs and products encoded by noncoding transcripts. In both species, the stress condition yielded fewer predicted translons overall, with pronounced reductions in uORFs and translons in noncoding transcripts, indicating class-dependent differences in the predicted response to stress. This atlas extends condition-resolved initiation predictions beyond the experimentally profiled transcripts and provides candidates for investigating sequence-dependent regulation of start-site selection.

### IncepTIS-guided design of stress-responsive translational elements

We next developed IncepTIS-evolve, a framework that uses condition-specific IncepTIS predictions to guide de novo sequence design. Starting from sequences generated using Evo 2, a genetic algorithm iteratively optimizes mutable regions while preserving specified fixed sequences (Fig. 6f; Methods and Supplementary Methods). We used this framework to design 282-nt 5′ leaders upstream of Renilla luciferase, keeping the luciferase Kozak sequence, start codon and coding sequence fixed. The optimization objective favored initiation at specified upstream sites under untreated conditions and at the luciferase start site under stress.

We independently designed three leaders with one, two or three target uORF initiation sites. The target positions were specified without fixing the corresponding start-codon sequences, allowing these sites to emerge during optimization. For each optimization, the highest-fitness sequence satisfying all success criteria was selected for experimental testing. IncepTIS predicted preferential initiation at the designed upstream sites under untreated conditions and increased initiation at the luciferase start site under stress (Fig. 6g).

To test these predictions, we introduced the designed leaders into dual-luciferase reporters and measured reporter activity in HEK293T cells with or without sodium arsenite treatment. Reporters containing the ATF4 5′ UTR and the parental vector were included for comparison. All three designed leaders showed an increase in the Renilla-to-firefly luciferase activity ratio after arsenite treatment, with the magnitude of the response varying among designs (Fig. 6h). These results support the use of condition-resolved initiation predictions to design synthetic 5′ leaders with experimentally measurable stress-responsive reporter output.

## DISCUSSION

Ribosome profiling has transformed our view of the coding capacity of mammalian transcriptomes by revealing widespread translation outside annotated coding sequences^5,6^. However, translon identification based on elongating-ribosome periodicity has difficulty resolving overlapping translation, particularly intORFs whose footprints are superimposed on those of the annotated CDS^16^. Although intORFs can be detected when their translation signals are sufficiently strong relative to those of the overlapping CDS^24^, identifying downstream initiation that generates N-terminally truncated variants remains particularly challenging with conventional Ribo-seq because these variants share the reading frame of the annotated CDS. Translation initiation profiling can improve start-site detection, but inhibitors such as harringtonine and lactimidomycin can perturb ribosome behavior and generate initiation-like signals from elongating ribosomes^28^. By selectively capturing eIF5B-associated ribosomes without harringtonine or lactimidomycin pretreatment, TISCA2 directly maps translation initiation sites and improves the detection of short and distinct translons, including intORFs, as well as N-terminally truncated variants.

This improved resolution revealed a broader landscape of stress-responsive translation initiation. The integrated stress response has traditionally been viewed largely through selective translation of uORF-regulated transcripts such as ATF4^36,45,46^. In contrast, TISCA2 showed that arsenite broadly redistributes start-site usage, reducing upstream initiation while increasing initiation downstream of annotated start codons. This shift increased the use of intORFs, truncated translons and other downstream translons that are difficult to resolve from elongating-ribosome profiles, with N-terminal proteomics supporting the production of selected downstream-initiated products. A more modest downstream shift was also observed in kidneys after LPS-induced systemic inflammation, extending these observations to an in vivo inflammatory setting. Thus, stress can remodel coding output not only by changing how much an mRNA is translated, but also by changing where translation begins. The mechanisms underlying this redistribution remain to be defined and may involve altered start-codon recognition, scanning and reinitiation.

Products of unannotated translons need not accumulate as stable proteins to have biological consequences^58,64,65^. Many predicted products of TISCA2-specific translons were short and showed low-confidence structural predictions, and a subset accumulated following proteasome inhibition, consistent with rapid proteostatic turnover.

Conversely, several MS-supported intORF-derived products exhibited distinct subcellular localization and interaction profiles, providing clues to their potential cellular roles. Rapid degradation may provide another route to biological relevance by supplying peptides for HLA-I presentation^58^, and our immunopeptidomic analysis identified HLA-I peptides derived from TISCA2-identified unannotated translons.

Notably, recent work has shown that unmutated unannotated translons, including intORFs, can generate cancer-restricted cryptic antigens that elicit cytotoxic T-cell responses and support tumor recognition^66^. Thus, unannotated translons can generate products with distinct cellular properties as well as transient products that remain biologically visible to the immune system.

TISCA2 revealed tissue-dependent differences in initiation-associated footprint distributions and candidate start-site usage. The testis showed greater relative footprint abundance downstream of annotated start codons and a larger contribution of downstream candidate TISs. Proteomic detection of selected predicted products provided additional support for translation at a subset of these sites. The testis also showed increased representation of AAG- and AGG-associated candidate TISs, although whether these signals represent genuine initiation remains uncertain.

Finally, the TISCA2 datasets enabled us to ask how much of start-site selection is encoded in mRNA sequence itself. Building on sequence-based approaches for predicting translation initiation sites and translational output^37–40^. IncepTIS predicts position-resolved initiation landscapes conditioned on cellular state. Its predictive performance on held-out transcripts suggests that mRNA sequence contains information relevant to condition-dependent start-site selection. Importantly, IncepTIS-guided optimization generated de novo 5′ leaders that conferred stress-responsive reporter output. This prospective design demonstrates that experimentally defined initiation landscapes can be translated into predictive and programmable sequence rules. Because the current model was trained primarily on HEK293T cells, transcriptome-wide predictions should nevertheless be regarded as a first-generation reference, and extending TISCA2 across additional tissues and physiological conditions should enable increasingly context-aware models of translation initiation.

## Methods

### Cell lines and culture conditions

HEK293T cells were obtained from the American Type Culture Collection (ATCC). Cells were cultured in Dulbecco’s modified Eagle’s medium (DMEM) supplemented with 10% fetal bovine serum (FBS), 100 U/mL penicillin–streptomycin (Fujifilm Wako), 1× non-essential amino acids (Fujifilm Wako), 2 mM L-glutamine (Fujifilm Wako), 1 mM sodium pyruvate (Fujifilm Wako), and 0.5 mM 2-mercaptoethanol (Fujifilm Wako). Cells were maintained at 37°C in a humidified atmosphere containing 5% CO_2_. All cell lines were tested for mycoplasma contamination using the MycoAlert Mycoplasma Detection Kit (Lonza) and were confirmed to be mycoplasma-free.

### Generation of eIF5B-3×FLAG knock-in HEK293T cells

Sense and antisense oligonucleotides encoding sgRNAs were cloned into pX330. Targeting constructs containing selection markers for homologous recombination were generated in pBluescript II SK (Stratagene) using the NEBuilder HiFi DNA Assembly Master Mix (New England Biolabs). HEK293T cells were transfected with the targeting vectors and sgRNA-expressing pX330 plasmids for 24 h and subsequently selected with puromycin for 2 weeks. Clonal cell lines were established by limiting dilution and screened by immunoblotting. Successful knock-in clones were further validated by genomic sequencing. eIF3D-V5 knock-in cells were established previously^29^.

### Cell treatments

For arsenite stress experiments used for Sel-TCP-seq and TI-seq, HEK293T cells were treated with 40 μM sodium arsenite for 1 h before harvesting. For whole-cell proteomic analysis, cells were treated with 40 μM sodium arsenite for 4 h, whereas cells used for HLA class I immunopeptidomics were treated with 40 μM sodium arsenite for 8 h. For experiments involving proteasome inhibition, cells were treated with 10 μM bortezomib (Btz) for 4 h before harvesting.

### Animals

All animal experiments were approved by the Animal Ethics Committees of Nagoya University (S230016, S240004, S250002 and S260003) and were conducted in accordance with institutional guidelines and regulations for animal care and use. To generate eIF5B-3×FLAG and eIF3D-V5 knock-in mice, CRISPR RNA (crRNA) and trans-activating CRISPR RNA (tracrRNA) were complexed with recombinant Cas9 (Integrated DNA Technologies) to form ribonucleoprotein complexes. C57BL/6J mouse zygotes were electroporated with the respective ribonucleoprotein complexes. Mice were maintained in a specific pathogen-free animal facility at Nagoya University under controlled conditions of 22°C, 50–60% humidity and a 12-h light/12-h dark cycle, with free access to water and CA-1 rodent chow (CLEA Japan).

### Chronic restraint stress

Mice were subjected to chronic restraint stress using a restraint cage (KN-469; Natsume Seisakusho) for 6 h per day for 14 consecutive days. Control mice were maintained under otherwise identical housing conditions without restraint. Following the final stress session, mice were subjected to tissue collection.

### LPS-induced systemic inflammation

Lipopolysaccharide (LPS; *Escherichia coli* O111:B4) was dissolved in PBS at 1 mg/ml and administered to mice by tail-vein injection at 5 mg/kg body weight. Control mice received an equivalent volume of PBS. Kidneys were collected 16 h after injection.

### Sel-TCP-seq in cultured cells

HEK293T cells were seeded at 5.0 × 10^6^ cells per 15-cm dish, using 12 dishes for each sample, and cultured for 2 days before harvesting. For arsenite-treated samples, cells were treated with 40 μM sodium arsenite for 1 h before crosslinking. Cells were incubated for 15 min at room temperature with crosslinking buffer containing PBS, 0.025% formaldehyde, 0.5 mM dithiobis(succinimidyl propionate) (DSP), 10 mM MgCl_2_ and 100 μg/ml cycloheximide. Crosslinking was quenched by addition of glycine. Cells were collected, washed with ice-cold PBS and lysed in buffer containing 50 mM HEPES-KOH (pH 7.5), 200 mM KCl, 10 mM MgCl_2_, 1% NP-40, 100 μg/ml cycloheximide and cOmplete EDTA-free protease inhibitor cocktail (Roche). Lysates were incubated for 10 min at 4°C and clarified by centrifugation at 20,380 × *g* for 10 min at 4°C. RNA concentration in the supernatant was measured using a NanoDrop spectrophotometer (Thermo Fisher Scientific). Lysates were digested with RNase I (100 U per 120 μg RNA; Thermo Fisher Scientific) for 5 min on ice and immediately loaded onto 15–50% sucrose gradients containing 20 mM Tris-HCl (pH 7.5), 160 mM KCl and 10 mM MgCl_2_. Gradients were centrifuged at 35,000 rpm (approximately 210,000 × *g*) for 5 h at 4°C in an SW41Ti rotor (Beckman Coulter) and fractionated using a gradient profiling system while monitoring absorbance at 260 nm. Fractions corresponding to 40S ribosomal complexes were pooled for immunoprecipitation. For Sel-TCP-seq, Protein G Dynabeads were preincubated with anti-FLAG or anti-V5 antibody in lysis buffer containing BSA overnight at 4°C. Pooled 40S fractions were incubated with antibody-conjugated beads for 90 min at 4°C. Beads were washed four times with lysis buffer before RNA purification.

### Sel-TCP-seq RNA purification and library preparation

Immunoprecipitated complexes were resuspended in 250 μl de-crosslinking buffer containing 1% SDS, 10 mM EDTA, 10 mM Tris-HCl (pH 7.5), 5 mM DTT and 10 mM glycine, followed by addition of 750 μl ISOGEN LS (Nippon Gene) and 200 μl chloroform. Samples were incubated for 45 min at 65°C and RNA was recovered by isopropanol precipitation. Purified RNA was size-selected over the range of 17–100 nt on a 15% polyacrylamide TBE–urea gel. RNA fragments were dephosphorylated, ligated to a DNA linker and depleted of rRNA using the riboPOOL Ribo-seq 96 reaction kit (human/mouse/rat; Biozym). Reverse transcription was performed using primer SI-019, and cDNA products were purified by denaturing PAGE. A second linker (SI-018) was ligated using T4 RNA Ligase 1 (New England Biolabs) overnight at 25°C. Libraries were amplified by PCR, size-selected on a 15% non-denaturing polyacrylamide gel and sequenced on a NovaSeq 6000 system (Illumina).

### Sel-TCP-seq in mouse tissues

Mice were anesthetized and subjected to transcardial perfusion with approximately 20 ml crosslinking buffer containing PBS, 0.5% formaldehyde, 10 mM MgCl_2_ and 100 μg/ml cycloheximide. Brain, kidney and testis were dissected immediately after perfusion and immersed in crosslinking buffer for an additional 20 min. Tissues were then transferred to quenching buffer containing glycine, MgCl_2_ and cycloheximide and incubated for 5 min at 4°C. Samples were transferred to tubes containing lysis buffer and snap-frozen in liquid nitrogen before storage at −80°C. Frozen tissues were mechanically disrupted by multi beads shocker (Yasui Kikai) in lysis buffer containing 50 mM Tris-HCl (pH 7.5), 150 mM NaCl, 5 mM MgCl_2_, 1% Triton X-100, 1 mM DTT, 100 μg/ml cycloheximide and cOmplete EDTA-free protease inhibitor cocktail. Lysates were clarified by centrifugation, and RNA concentration in the supernatant was determined using a NanoDrop spectrophotometer. Lysates were digested with RNase I (100 U per 120 μg RNA) for 5 min on ice and immediately loaded onto 15–50% sucrose gradients containing 20 mM Tris-HCl (pH 7.5), 150 mM NaCl, 5 mM MgCl_2_ and 1 mM DTT. Gradients were centrifuged at 35,000 rpm for 5 h at 4°C in an SW41Ti rotor and fractionated while monitoring absorbance at 260 nm. Fractions corresponding to 40S ribosomal complexes were pooled and subjected to immunoprecipitation using anti-FLAG or anti-V5 antibody-conjugated Protein G Dynabeads. RNA purification and library preparation were performed as described above for cultured cells.

### RNA-seq

Total RNA was extracted from cell lysates or tissue with the use of a PureLinkRNAMini Kit (Thermo Fisher Scientific), and the quality of the purified RNA was assessed with a 2100 Bioanalyzer (Agilent). After mRNA selection with the use of a NEBNext Poly(A) mRNA Magnetic Isolation Module (New England BioLabs), libraries were prepared with the use of a NEB Next Ultra Directional RNA Library PrepKit for Illumina (New England BioLabs). The cDNAs were sequenced with a NovaSeq 6000 system.

### Ribo-seq and TI-seq

For TI-seq, HEK293T cells were treated with 2 μg/ml harringtonine (MedChemExpress) for 3 min at 37°C before harvesting. Cells were immediately placed on ice, washed with ice-cold PBS and lysed in buffer containing 50 mM Tris-HCl (pH 7.5), 150 mM NaCl, 5 mM MgCl_2_, 1 mM DTT, 1% Triton X-100 and EDTA-free protease inhibitor cocktail, supplemented with either 100 μg/ml cycloheximide for Ribo-seq or 2 μg/ml harringtonine for TI-seq. Lysates were treated with TURBO DNase (Thermo Fisher Scientific) for 10 min on ice and clarified by centrifugation at 20,380 × *g* for 10 min at 4°C. RNA concentration was measured using the Qubit RNA BR Assay Kit (Thermo Fisher Scientific), and lysates were digested with RNase I (20 U per 10 μg RNA; Epicentre) for 45 min at 25°C. Digestion was terminated by addition of SUPERase•In RNase inhibitor (Invitrogen). Samples were layered onto a 1 M sucrose cushion containing 20 mM Tris-HCl (pH 7.5), 150 mM NaCl, 5 mM MgCl_2_, 1 mM DTT, 100 μg/ml cycloheximide and 10 U/ml SUPERase•In, and centrifuged at 100,000 rpm (417,200 × *g*) for 1 h at 4°C in a TLA110 rotor (Beckman Coulter). Ribosome-protected RNA fragments were purified using ISOGEN (Fujifilm) and size-selected on a 15% polyacrylamide TBE–urea gel. Fragments of 17–34 nt were recovered and treated with T4 polynucleotide kinase (New England Biolabs), followed by ligation of barcoded DNA linkers (NI-810 to NI-817) using T4 RNA Ligase 2, truncated K227Q (New England Biolabs). After gel purification, rRNA was depleted using the riboPOOL Ribo-seq 96 reaction kit (human/mouse/rat; Biozym). Reverse transcription was performed using primer NI-802, and the resulting cDNAs were circularized using CircLigase II (Lucigen). Libraries were amplified by PCR using NI-798 together with indexed primers NI-799 and NI-822 to NI-826, size-selected on a 15% non-denaturing polyacrylamide gel and sequenced on a NovaSeq 6000 system (Illumina).

### Nanopore long-read RNA sequencing

Full-length cDNA was prepared from 1 µg of total RNA using a modified protocol of full-length cDNA-selective sequencing (FLeCS-seq). The original FLeCS-seq selectively synthesizes complete full-length cDNA by capturing both the 5’ cap structure with improved oligo-capping^67^ and the polyA tail with splint ligation. In the modified protocol, oligo-dT priming was employed for 3’ end capture instead of polyA ligation.

Specifically, the polyA ligation step was omitted, and the improved oligo-capping was performed directly on the denatured total RNA. For first-strand cDNA synthesis, an oligo-dT primer (AAGCAGTGGTATCAACGCAGAGTACTTTTTTTTTTTTTTTTTTTTTTTTTTTTT TVN) was used in the annealing step prior to reverse transcription. PCR amplification was performed under the following conditions: initial denaturation at 94°C for 1 min; 10 cycles of 94°C for 15 s, 63°C for 5 s, and 68°C for 15 min; final extension at 68°C for 15 min; hold at 4°C. Nanopore sequencing libraries were prepared using the Ligation Sequencing Kit V14 (SQK-LSK114, Oxford Nanopore Technologies). Each library was sequenced using a PromethION flow cell (FLO-PRO114M; Oxford Nanopore Technologies) and the PromethION sequencer. Base calling was performed using Dorado 0.8.2 with the dna_r10.4.1_e8.2_400bps_sup@v5.0.0 model. Raw reads were trimmed of adapter sequence using Pychopper v2.4.0, and the trimmed reads with polyA sequence on the 3′ ends were selected as in our previous study. The reads were aligned to the reference genome hg38 or mm10 for human and mouse samples, respectively, using minimap2 v2.28^68^. RNA isoforms were detected using StringTie v3.0.1^69^ guided by GENCODE v48 and vM25 for human and mouse samples, respectively.

### Processing of Sel-TCP-seq, TI-seq, Ribo-seq and RNA-seq data

Adaptor sequences were trimmed from raw reads using cutadapt. Low-quality reads were discarded using the fastq_quality_trimmer and fastq_quality_filter tools in the FASTX Toolkit. Ribosomal RNA reads were removed by alignment with human rRNA sequences using STAR, and the remaining reads were aligned to the human or mouse transcriptome annotated by StringTie and genome (hg38 or mm10) using this software. Multiple mappings were performed. The metagene analysis was performed using custom Python scripts in Numpy (v1.17.3) and Pysam (v0.15.3). The P-site was defined as the position located 12 nt from the start of the 21-, 28- and 29-nt reads, and 13 nt from the start of the 22-nt reads, that showed a 3-nt periodicity. Three-nucleotide periodicity was analyzed using RibORF, and footprints with a score of ≥0.5 were applied to the analysis. RNA-seq reads mapped to human or mouse coding transcripts were counted using featureCounts (the Subread package) and transcripts per kilobase million (TPM) values were calculated. P-site corrected Ribo-seq reads were counted using a custom Python script. Translation efficiency (TE) was calculated for each translon according to the formula: TE = TPM for translon in Ribo-seq/TPM for transcript in RNA-seq. eIF3D Sel-TCP-seq, Ribo-seq, TI-seq and RNA-seq data sets were retrieved from GEO [GSE174329 and GSE308359]^29,70^. To calculate precision and recall, TI-seq data were retrieved from GEO [GSE174329]. The mapped read data were converted into read aggregation plots by Numpy (v1.17.3) and Pysam (v0.15.3). After normalization by the total number of reads, the points at which the derivative of TI-seq normalized footprint density exceeded the threshold were defined as TI-seq peaks for each transcript. Where μ is the mean, σ is the standard deviation, and n is an integer from 1 to 20, μ + nσ was used as the threshold. Among peaks accompanied by AUG or near-cognate codons on the downstream side, those with a known start codon at a position +12 to +14 nt from the peak were defined as positive. ORF-RATER was run using Ribo-seq and TI-seq data from HEK293T cells with or without arsenite treatment.

### TISCA2 identification of translation initiation sites

We first generated a transcript isoform catalog from full-length cDNA sequencing and retained expressed mRNA isoforms with at least ten long-read coverage and ten eIF5B Sel-TCP-seq read counts. For these transcripts, all AUG codons and near-cognate codons differing from AUG by one nucleotide were considered candidate TISs.

Candidate sites were then filtered by length coverage, defined as the number of distinct footprint lengths represented within a 60-nt window around the candidate site. Because canonical TISs with length coverage of at least ten showed substantially higher cosine similarity, this threshold was used to remove poorly covered candidates (Extended Data Fig. 2a–c). For each remaining candidate TIS, TISCA2 calculated cosine similarity to the eIF5B differential metagene signature. Using annotated AUG start codons of expressed coding transcripts as positives, we selected a cosine-similarity cutoff that balanced precision and recall (Extended Data Fig. 2d,e). Candidates exceeding this cutoff and forming local maxima were called as TISCA2-positive TISs. In this benchmark, the eIF5B-based TISCA2 score outperformed inhibitor-based TI-seq in precision-recall analysis (Extended Data Fig. 2d), supporting the utility of initiating-ribosome capture for physiological TIS detection. We used a classification adapted from Chen et al. for grouped comparisons^53^. For grouped comparisons, annotated translons corresponded to annotated CDSs; variant translons comprised extended and truncated ORFs; and distinct translons comprised uORFs, uoORFs, intORFs, doORFs and dORFs. Translons classified as noncoding or novel were analyzed separately and were excluded from comparisons between annotated/variant and distinct translons. In the translon classification, "novel" refers to translons on novel transcripts identified by StringTie (with IDs beginning with “MSTRG”). Selenoproteins were classified as intORFs. For TISs with overlapping genomic coordinates, only those with the highest long-read coverage and translon priority order were retained. The translon priority order was as follows: annotated > extended > uORF > uoORF > truncated > intORF > dORF > doORF > novel > noncoding.

### Calculation of initiation efficiency

To reduce background, we defined effective reads as footprints whose 3′ ends mapped between +15 and +30 nt relative to the candidate start codon within the 60-nt TIS window (Extended Data Fig. 2g). We then calculated initiation efficiency (IE) as eIF5B effective reads per million (RPM) divided by RNA abundance measured by RNA-seq transcripts per million (TPM).

### Leaky-scanning score

For all coding transcripts with a 5’ UTR longer than 50 nt, reads mapping from −50 nt to +30 nt relative to the start codon were counted as the amount of initiating ribosomes, and reads mapping downstream of +30 nt from the start codon toward the 3’ UTR were counted as the amount of leaky scanning ribosomes. The leaky-scanning score was calculated as the log_2_ ratio of downstream footprint counts to start-codon-proximal footprint counts. Transcripts were excluded if either the initiating ribosome read count or the leaky scanning ribosome read count was less than 10.

### Sample preparation for N-terminal proteomics using HUNTER

SP3 beads were prepared by mixing hydrophobic Sera-Mag SpeedBeads (catalog no. 6515-2105-050250; Thermo Fisher Scientific) and hydrophilic Sera-Mag SpeedBeads (catalog no. 4515-2105-050250; Thermo Fisher Scientific) at a 1:1 ratio. The resulting SP3 beads were washed with water and resuspended in Milli-Q water. Cells were lysed in buffer containing 1% SDS, 50 mM Tris-HCl (pH 7.5), and 150 mM NaCl, and the lysates were immediately heated for 5 min. Brain, kidney, and testis tissues were collected from 10-week-old male mice and homogenized in the same buffer using a Multi-beads Shocker (YASUI KIKAI). The tissue homogenates were immediately heated for 5 min. Cell lysates and tissue homogenates were then treated with Benzonase nuclease (Merck, catalog no. 71205-3CN) at 37°C for 15 min and centrifuged at 20,000 × g for 10 min at 4°C. The supernatants were collected, and protein concentrations were determined using a BCA assay. Protein aliquots (50 µg) were reduced with 2.5 mM TCEP at 37°C for 30 min and alkylated with 12.5 mM iodoacetamide at room temperature for 30 min, followed by SP3-based protein cleanup. Briefly, 2 µL of SP3 bead suspension was added to each sample, and ethanol was added to a final concentration of 80%. After mixing for 30 min, the beads were washed twice with 90% ethanol, and the bead-bound proteins were resuspended in 100 µL of 100 mM HEPES (pH 7.4). Primary amino groups were then dimethylated by adding formaldehyde and sodium cyanoborohydride (catalog no. S0395; Tokyo Chemical Industry) to final concentrations of 30 and 15 mM, respectively, followed by incubation at 37°C for 2 h. The same amounts of both reagents were added again, and the reaction was continued for an additional 2 h. The reaction was quenched with 10 µL of 1.0 M Tris-HCl (pH 6.8) for 1 h. Proteins were rebound to SP3 beads by adding 440 µL of ethanol and 1 µL of SP3 bead suspension, mixed for 30 min, and washed twice with 80% ethanol. The beads were resuspended in 50 µL of 100 mM HEPES (pH 8.0) and digested with 2 µg of trypsin for 16 h. The peptide-containing supernatant was recovered, heated at 95°C for 5 min, adjusted to 40% ethanol, and reacted with undecanal (catalog no. U0009; Tokyo Chemical Industry) at an undecanal-to-peptide ratio of 50:1 (w/w). After addition of 1 µL of 100 mM HCl, sodium cyanoborohydride was added to a final concentration of 10 mM, and the mixture was incubated at 37°C for 1 h. An additional equal amount of sodium cyanoborohydride was then added, followed by incubation at 37°C for 1 h. The reaction mixture was acidified to 0.5% trifluoroacetic acid while maintaining 40% ethanol and loaded onto an SDB-Stage tip preconditioned with methanol and equilibrated with 0.1% trifluoroacetic acid in 40% ethanol. A peptide amount corresponding to 10 µg of protein was loaded onto the SDB-Stage tip.

Undecanal-modified peptides were retained on the SDB-Stage tip, whereas the flow-through fraction containing acetylated or dimethylated N-terminal peptides was collected and dried using a vacuum concentrator.

### Sample preparation for total proteomics

HEK293T cells were seeded the day before harvest. Four hours before collection, sodium arsenite (final concentration, 40 μM) and/or bortezomib (Btz; final concentration, 10 μM; FUJIFILM Wako) were added to the culture medium as indicated. Cells were then washed once with PBS and lysed in lysis buffer [20 mM Tris–HCl (pH 7.5), 150 mM NaCl, 1% Triton X-100, and protease inhibitor cocktail (cOmplete, Roche)]. Lysates were incubated on ice for 10 min and clarified by centrifugation at 20,000 × *g* for 10 min at 4°C, after which the supernatants were collected. Brain, kidney, and testis tissues were collected from 10-week-old male mice and homogenized in the same lysis buffer using a Multi-beads Shocker (YASUI KIKAI). The homogenates were centrifuged at 20,000 × *g* for 10 min at 4°C, and the supernatants were collected. Protein concentrations were determined using a BCA Protein Assay Kit (BioDynamics Laboratory Inc.), and 10 μg of protein from each sample was used for subsequent analyses. Cysteine residues were reduced with 2.5 mM tris(2-carboxyethyl)phosphine hydrochloride (Thermo Fisher Scientific) for 30 min at 37°C and alkylated with 50 mM 2-iodoacetamide (Merck) for 30 min at room temperature. Samples were purified and digested using the SP3 method. The washed beads were resuspended in 100 μl of 50 mM Tris–HCl (pH 8.0) containing 0.2 μg of Lys-C (FUJIFILM Wako) and incubated at 37°C for 4 h. Subsequently, 0.2 μg of trypsin (Promega) was added, and the samples were incubated overnight at 37°C with gentle mixing. After digestion, the samples were placed on a magnetic rack to separate the beads. The supernatants were collected, snap-frozen in liquid nitrogen, and stored at −80°C until further analysis.

### Sample preparation for HLA-I immunopeptidomics

1 × 10⁹ HEK293T cells were used to isolate HLA-bound peptides, as previously described^71^. The sample was ground under cryogenic conditions and lysed with a lysis buffer containing 0.25% sodium deoxycholate, 0.2 mM iodoacetamide, 1 mM EDTA, a protease inhibitor cocktail (Sigma), 1 mM PMSF, and 1% octyl-β-D-glucopyranoside (Dojindo) in DPBS. HLA-I-bound peptides were captured as peptide–HLA complexes using affinity chromatography with the W6/32 mAb (HB-95, ATCC), which recognizes pan-HLA-I, coupled to CNBr-activated Sepharose 4B (GE Healthcare) overnight. The peptides were then eluted with 0.2% TFA and desalted using Sep-Pak tC18 (Waters) with 28% ACN in 0.1% TFA, followed by ZipTip U-C18 (Millipore) with 50% ACN in 1% FA. The eluate was dried by vacuum centrifugation. Hybridomas producing the mAb were cultured in hybridoma serum-free medium (SFM; Gibco) supplemented with 1% penicillin–streptomycin in CELLine Bioreactor Flasks (CL1000, Corning). The concentrated mAb was collected and purified using a HiTrap Protein G HP column (GE Healthcare).

### LC–MS/MS analysis

HUNTER-enriched peptides were analyzed using an Evosep LC system coupled to an Orbitrap Astral mass spectrometer equipped with an EASY-Spray ion source (Thermo Fisher Scientific). Peptides were separated on an EASY-Spray PepMap RSLC C18 column (ES906; 150 mm × 150 μm i.d., 2 μm particles) maintained at 45 °C using the 30 SPD method with a 44-min gradient. MS acquisition was performed for 46 min in positive-ion data-independent acquisition (DIA) mode without FAIMS. MS1 spectra were acquired in the Orbitrap analyzer at a resolution setting of 240,000 over m/z 300–1500, with a normalized AGC target of 500% and a maximum injection time of 20 ms. DIA acquisition was configured with 240 non-overlapping isolation windows of nominal width 5 Th across a precursor range of m/z 300–1500, with window placement optimization enabled. Higher-energy collisional dissociation (HCD) was performed at a normalized collision energy of 27%. MS/MS spectra were acquired in the Astral analyzer over m/z 150–2000, with a normalized AGC target of 800% and a maximum injection time of 5 ms. Differential analysis was performed using Perseus^72^.

For total proteome analysis, peptides were analyzed using an Evosep One LC system coupled to an Orbitrap Astral mass spectrometer (Thermo Fisher Scientific). Peptides were separated on an Evosep C18 column (EV1137) using the 30 SPD method with a 44-min gradient. MS acquisition was performed in data-independent acquisition (DIA) mode. MS1 spectra were acquired in the Orbitrap analyzer at a resolution of 240,000 over m/z 380–980, with an AGC target of 500% and a maximum injection time of 5 ms. DIA MS/MS spectra were acquired in the Astral analyzer using 2 Th isolation windows, an HCD collision energy of 25%, an AGC target of 500%, and a maximum injection time of 3 ms. All DIA raw data were processed with DIA-NN (version 2.3.2) in the library-free search mode with reference to human or mouse UniProt sequences and TISCA2- or ORF-RATER-derived databases.

Immunopeptides were analyzed using the same LC–MS platform and column at 45 °C with the Evosep 15 SPD Extended method. MS acquisition was performed for 88 min in positive-ion data-dependent acquisition (DDA) mode without FAIMS. MS1 spectra were acquired in the Orbitrap analyzer at a resolution setting of 120,000 over m/z 300–1500, with a normalized AGC target of 300% and a maximum injection time of 100 ms. Precursors with charge states of 1–4 and a minimum intensity of 5,000 were selected using a 2 Th isolation window and fragmented at a normalized HCD collision energy of 30%. MS/MS spectra were acquired in the Astral analyzer over m/z 100–1000 using the standard AGC setting and a maximum injection time of 10 ms. The DDA cycle time was set to 0.6 s, with dynamic exclusion of 15 s. Raw MS data were analyzed using MSFragger (version 4.4.1) within FragPipe (version 23.1). MS/MS spectra were searched against TISCA2- or ORF-RATER-derived databases using a non-specific digestion setting without enzyme cleavage constraints. Peptide lengths were restricted to 7–25 amino acids, and precursor charge states of 1–4 were considered. The precursor mass tolerance was set to 20 ppm, and the fragment mass tolerance was initially set to 20 ppm and subsequently optimized by MSFragger. Carbamidomethylation of cysteine was not specified as a fixed modification. Methionine oxidation, protein N-terminal acetylation, N-terminal glutamine and glutamic acid cyclization, and cysteine modification (+119.0041 Da) were allowed as variable modifications, with a maximum of three variable modifications per peptide. Protein N-terminal methionine excision was enabled. Peptide-spectrum matches (PSMs) were further evaluated using MSBooster (version 1.3.17) with DIA-NN-based predictions of fragment ion spectra and retention times, followed by rescoring using Percolator (version 3.7.1). Protein inference was performed using ProteinProphet, and identifications were filtered using Philosopher (version 5.1.2) with a target-decoy strategy to control the false discovery rate (FDR) at 1% at the PSM and peptide levels.

### NetMHCpan analysis

NetMHCpan scores for 8–12 mer peptides were calculated for all translon sequences predicted by TISCA2 or ORF-RATER. For strong binding peptides, the highest score among those for the HLA-A02:01, HLA-A03:01, HLA-B07:02, HLA-B35:08, and HLA-C07:02 alleles was adopted.

### Structural prediction using AF3

Atomic models of all selected translon products were predicted using AlphaFold3^56^. Structure prediction was performed with all first amino acids substituted with methionine.

### ESM-2 sequence embedding

Protein sequence embeddings were generated using the pretrained ESM-2 model (ESM2_t33_650M_UR50D), which consists of 33 Transformer layers. For each amino acid sequence, residue-level representations were extracted from the final (33rd) layer, and the representations were averaged across all residues to obtain a single sequence-level embedding vector.

### Cloning of intORF products

Each internal ORF was amplified from HEK293T cell cDNA and cloned into pcDNA3.1 using the XbaI and BamHI restriction sites to generate constructs encoding N-terminally V5-tagged intORF products.

### Immunoblot analysis

Protein samples were subjected to sodium dodecyl sulfate-polyacrylamide gel electrophoresis on SuperSep Ace 5-20% (Fujifilm). Membranes were incubated consecutively with primary antibodies [V5 tag (Thermo Fisher Scientific, R960CUS), eIF2α (Cell Signaling Technology, 5324S), Phospho-eIF2α (ser51) (Proteintech, 68023-1-IG) and HSP90 (BD Biosciences, 610419)] and horseradish peroxidase-conjugated secondary antibodies (Promega), and signals were visualized using SuperSignal West (Thermo Fisher Scientific) reagents on a ChemiDoc imaging system (Bio-Rad).

### RT-qPCR analysis

Total RNA was extracted from kidney tissues using the PureLink RNA Mini Kit (Thermo Fisher Scientific) and reverse-transcribed using ReverTra Ace qPCR RT Master Mix (TOYOBO). Quantitative real-time PCR was performed using the resulting cDNA, gene-specific primers, and SYBR Green qPCR Master Mix (Thermo Fisher Scientific) on a QuantStudio 3 Real-Time PCR System (Applied Biosystems). Target mRNA levels were normalized to β-actin mRNA levels.

### Immunofluorescence microscopy

HEK293T cells were transfected with pcDNA3-based expression plasmids encoding N-terminally V5-tagged proteins using X-tremeGENE 9 (Roche). Forty-eight hours after transfection, cells were processed for immunofluorescence analysis. Cells were washed once with PBS and fixed with 4% paraformaldehyde in PBS for 10 min at room temperature. After fixation, cells were washed three times with PBS and permeabilized and blocked in staining buffer (PBS containing 1% bovine serum albumin, 0.3% Triton X-100, and 0.01% sodium azide) for 1 h at room temperature. Coverslips were then incubated with anti-V5 primary antibody diluted in staining buffer overnight at 4°C. Following three washes with staining buffer, coverslips were incubated with Alexa Fluor 488–conjugated secondary antibody for 1 h at room temperature in the dark. After three additional washes, coverslips were mounted using VECTASHIELD Hard Set Mounting Medium with DAPI (Vector Laboratories) and imaged on a Zeiss LSM710 confocal microscope.

### V5 immunoprecipitation and interactome mass spectrometry

HEK293T cells were transfected with pcDNA3-based expression plasmids encoding N-terminally V5-tagged proteins using X-tremeGENE 9 (Roche). Forty-eight hours after transfection, cells were washed once with PBS and lysed in lysis buffer [50 mM Tris–HCl (pH 7.5), 150 mM NaCl, 1% Triton X-100, 1 mM PMSF, and protease inhibitor cocktail (cOmplete, Roche)]. Lysates were incubated on ice for 10 min and clarified by centrifugation at 20,000 × g for 10 min at 4°C. The supernatants were collected, and protein concentrations were determined using a BCA assay. For immunoprecipitation, Dynabeads protein G (VERITAS) were incubated overnight with an anti-V5 antibody (Invitrogen) in the presence of 0.5 mg/mL BSA. The antibody-coated beads were then incubated with lysates containing 1.5 mg of total protein for 3 h at 4°C with rotation.

The beads were washed three times with lysis buffer, and bound proteins were eluted by incubation in lysis buffer containing 0.5 mg/mL V5 peptide (Merck) for 10 min at room temperature. The purified proteins were precipitated with trichloroacetic acid and washed once with 100% acetone and twice with 90% acetone and resuspended in a buffer containing 4% SDS, 100 mM Tris–HCl, and 20 mM NaCl. The immunoprecipitates were incubated for 15 min at 37°C in 25 mM tris(2-carboxyethyl)phosphine hydrochloride and alkylated for 30 min at 37°C in the dark using 25 mM iodoacetamide. The reduced and alkylated proteins were processed using the single-pot, solid-phase-enhanced sample preparation (SP3) method. Proteins were digested on the SP3 beads with Trypsin/Lys-C Mix (Promega) overnight at 37°C. Digested peptides were desalted using GL-Tip SDB (GL Sciences, Tokyo, Japan), evaporated in a SpeedVac concentrator, and redissolved in 0.1% trifluoroacetic acid and 2% acetonitrile. Peptides were analyzed by nano-flow liquid chromatography–tandem mass spectrometry (nanoLC–MS/MS) using a Q Exactive Orbitrap mass spectrometer (Thermo Fisher Scientific) coupled to a nanoelectrospray ionization source (AMR Inc., Tokyo, Japan). Peptides were separated on a C18 reversed-phase column (100 μm inner diameter × 125 mm length; Nikkyo Technos, Tokyo, Japan). The mobile phases consisted of 0.5% acetic acid in water (A) and 80% acetonitrile containing 0.5% acetic acid (B). Peptides were separated using a linear gradient of 5–40% B over 100 min, followed by an increase to 95% B over 5 min at a flow rate of 300 nL/min. The mass spectrometer was operated in data-independent acquisition (DIA) mode. Full MS spectra were acquired over an m/z range of 500–740 at a resolution of 70,000, with an automatic gain control (AGC) target of 3 × 10⁶ and a maximum injection time of 10 ms.

DIA MS/MS spectra were acquired using 24 isolation windows of 10 m/z each at a resolution of 70,000, with an AGC target of 3 × 10⁶ and automatic maximum injection time. The normalized collision energy (NCE) was set to 22, and the fixed first mass was set to m/z 200. All DIA raw data were processed with DIA-NN (version 2.3.2) in the library-free search mode with reference to human UniProt sequences and the bait proteins. Differential analysis was performed using Perseus^72^.

### Variant calling and phasing analysis

FASTQ files obtained from nanopore PromethION (Oxford Nanopore Technologies) were filtered for mapping quality ≧ 9 and length ≧ 200 base, using NanoFilt (2.8.0), and mapping to GRCh38 genomic sequence was conducted using minimap2 (2.26-r1175)^68^. Resulting mapping files were used for assembly using StringTie3 (3.0.0) and variant calling with Clair3_RNA (0.2.2)^69,73^. Then, called variants were annotated for haplotype phases using WhatsHap (2.8)^74^, and separated into two phases. Coordinates of GRCh38 genomic sequence and assembled transcriptional annotations were adjusted adhering variant positions for each phase, using bcftools (1.23.1) and CrossMap (0.7.0)^75^. Subsequently, transcriptional sequences with variant information of individual genomic phases were extracted using GenomicFeatures (1.58.0) and formatted into FASTA^76^.

### IncepTIS and IncepTIS-evolve

#### Dataset preparation

The training dataset comprised 62,106 samples derived from 15,531 transcripts profiled by TISCA2 in HEK293T, covering unstressed and arsenite-treated conditions with two biological replicates per condition. For the 16,896 samples in which the Nanopore long-read sequence matched the length of the corresponding reference sequence, the long-read sequence was used; for the remaining 45,210 samples, in which no long-read sequence was available or the lengths did not match, the Ensembl reference sequence was used. The TISCA2 cosine similarity profile was used as the continuous target at each position without normalization. Missing values were masked and, together with positions having negative cosine similarity, were excluded from both the loss and the evaluation. Positions with cosine similarity ≥ 0.2 were defined as positive and used both as the binary target and for PR-AUC evaluation (Supplementary Methods). The data were split into ten folds at the level of transcript identifier. Files derived from the same transcript across conditions and replicates were assigned to the same fold, so that no transcript was shared between the training and validation sets.

#### Model architecture

IncepTIS uses NTv3 (‘InstaDeepAI/NTv3_100M_pr’) as an RNA sequence encoder that extracts a per-nucleotide embedding^63^.

A 768-dimensional feature vector is obtained for each nucleotide position from the final layer. The condition was encoded,with 0 for the unstressed condition and 1 for the arsenite-stress condition, broadcast across all sequence positions and concatenated with the NTv3 features, as the input to the Attention U-Net prediction head.

The Attention U-Net prediction head consists of three downsampling convolutional blocks and three upsampling convolutional blocks, with an attention gate on each of the three skip connections. The model has 108M parameters in total, of which 1.73M belong to the Attention U-Net. The final layer of the Attention U-Net head applies a 1 × 1 convolution and predicts one logit per nucleotide. The regression output is obtained by applying a sigmoid to this logit, and the binary logit is obtained from the same pre-sigmoid logit by subtracting a learnable threshold parameter *τ* in logit space.

#### Loss function and model training

The loss was defined as a weighted sum of a binary cross-entropy (BCE) loss and a mean squared error (MSE) loss.

All parameters, including those of the NTv3 backbone, were fine-tuned together with the Attention U-Net. The model was optimized using Adam^77^ with an initial learning rate of 1 × 10⁻³, and training was performed by distributed data-parallel training on four GPUs. The learning rate was scheduled using the update-based cosine scheduler in timm^78^. Each fold was trained for 20 epochs, and early stopping was not used.

#### Model evaluation

The precision–recall area under the curve (PR-AUC) was used as the primary performance metric. PR-AUC was calculated over all valid nucleotide positions of the validation transcripts in each fold, and additionally from a single curve pooling the held-out predictions of all ten folds.

#### Transcriptome-wide prediction

For the transcriptome-wide atlas, predictions from the ten cross-validation models were averaged and quantized to 8-bit integers spanning the cosine-similarity range 0–1.

Positions with a value of at least 13 (≈0.05, lower than the 0.2 used for evaluation to favour sensitivity) were called positive.

#### Sequence optimization using IncepTIS-evolve

IncepTIS-evolve is a genetic algorithm that searches for sequences carrying an arbitrary set of initiation sites, using the per-nucleotide TISCA2 cosine similarity predicted by IncepTIS under the unstressed and stress conditions (Supplementary Algorithms 1 and 2).

The designed sequence length, start positions and fixed sequences of the target sites were specified. Nucleotides within the fixed segments were held constant throughout the search, and all other positions were treated as mutable. A subset of the initial population was generated with Evo 2 and overwritten at the fixed segments; the remaining sequences were generated by sampling mutable positions uniformly from A, C, G and T (Supplementary Methods).

In each generation, a fixed number of elite sequences with the highest fitness were carried over unchanged (Supplementary Methods). The remainder were generated by selecting two parents by tournament selection, applying uniform crossover, and then introducing mutations.

Independently of fitness, success was defined as the simultaneous satisfaction of a set of hard constraints on the predicted initiation signals at the target sites (Supplementary Methods). Among the sequences meeting all of these criteria, the one with the highest fitness was returned; only when no sequence met all criteria was the sequence with the highest fitness among all candidates returned instead.

### Luciferase reporter assay

Plasmids were generated by GenScript and transfected into cells using the X-tremeGENE 9 DNA Transfection Reagent (Merck). Cells were harvested after 48 h of incubation, and luciferase activity was measured using the Dual-Luciferase Reporter Assay System (Promega) on a GloMax Explorer Multimode Microplate Reader (Promega). HEK293T cells were treated with or without 40 μM sodium arsenite for 8 h before harvesting.

### Statistical analysis

The generalized linear model in RiboDiff was used to analyze the differential initiation efficiency^79^. Two-sided Student’s *t*-test, Mann–Whitney *U* tests and Hotelling’s *T*-squared test were performed using Scipy (v1.2.1) in Python 3.6.8. An adjusted *P*-value (false discovery rate q value < 0.05) was used for the extraction of differentially expressed genes. *P*-values <0.05 were otherwise considered statistically significant. GO analysis and EASE score calculation were performed in DAVID functional annotation bioinformatics microarray analysis.

### Code availability

The TISCA2 analysis pipeline and example data are available at https://github.com/kazu-ichihara/TISCA2. Code for variant calling and haplotype phasing is available at https://github.com/ajiromasa/variant_calling. The source code and software packages for IncepTIS and IncepTIS-evolve are available at https://github.com/HaLU-9000/IncepTIS.

## Supporting information

Supplementary Methods, Extended Data Figures, Supplementary Tables and Algorithms

## Acknowledgements

We thank all members of the Matsumoto Lab for helpful discussions. Computations were partially performed on the NIG supercomputer at ROIS National Institute of Genetics, the TSUBAME4.0 supercomputer at Institute of Science Tokyo, the supercomputer “Flow” at the Information Technology Center, Nagoya University, and the SHIROKANE supercomputer at the Human Genome Center, the University of Tokyo. This work was supported in part by MEXT/JSPS KAKENHI (Grant Numbers JP26K23165 to M.S., JP26K18235 to K.I. and JP20H05928 to A.M.); the Japan Science and Technology Agency (JST) FOREST Program (Grant Numbers JPMJFR2118 to A.Y. and JPMJFR2312 to A.M.); the Japan Agency for Medical Research and Development (AMED) (Grant Numbers JP26tm0424235 to M.S., JP26ama221315 to T.K., JP26ama221356 to S.T., JP22gm6410018, JP24tm0424231 and JP25wm0625327 to A.M.); the Foundation of Kinoshita Memorial Enterprise (Basic Science Fund, to A.M.); and the Mochida Memorial Foundation for Medical and Pharmaceutical Research (to K.I.). This publication has emanated from research conducted with the financial support of Taighde Éireann – Research Ireland under Grant number 25/FFP-A/13662 (Frontiers for the Future Programme, to P.V.B.).

## Author contributions

Conceptualization: A.M.; Methodology: K.I. and H.M.; Software: K.I. and H.M.; Formal analysis: K.I., H.M., S.S. and M.A.; Investigation: K.I., T. Shiraishi, C.S., M.H., N.Y., A.H., S.T., E.M.-S., A.S., K.A., M.S. and S.A.; Visualization: A.M., K.I., H.M. and T. Shimamura; Supervision: K.T., T.I., Y.S., A.Y., T.K., M.M., T. Shimamura and A.M.; Funding acquisition: M.S., A.Y., T.K., S.T., A.M., K.I. and P.V.B.; Writing – original draft: A.M., K.I., T. Shiraishi, H.M. and T. Shimamura; Writing – review & editing: A.M., K.I., T. Shiraishi, H.M., T. Shimamura and P.V.B.

## Competing interests

A.Y. has received research funding from Daiichi Sankyo, Chugai Pharmaceutical, Eisai, and xFOREST Therapeutics, unrelated to this study; serves as a scientific advisor to xFOREST Therapeutics; and has received honoraria from Chordia Therapeutics, Otsuka Pharmaceutical, MSD, Kyowa Kirin, Daiichi Sankyo, Sanofi, Sumitomo Dainippon Pharma, Life Technologies Japan, Amgen, Bayer, and Oxford Nanopore Technologies. P.V.B. is a co-founder and shareholder of EIRNABio Ltd. All other authors declare no competing interests.

**Correspondence** and requests for materials should be addressed to Teppei Shimamura or Akinobu Matsumoto.

