## Supplementary Methods, Extended Data Figures, Supplementary Tables and Algorithms for "Physiological mapping and prediction reveal stress-responsive translation initiation landscapes"

**Extended Data Figures**

**Supplementary Tables 1-3**

**Supplementary Algorithms 1 and 2**

### Supplementary Methods

#### IncepTIS Training Dataset Details

The Nanopore long-read data contained transcripts with identifiers beginning with MSTRG, but only transcripts carrying an Ensembl identifier were used in IncepTIS training.
Of the 15,531 transcripts in the training dataset, 2 replicates were obtained for every transcript under the unstressed condition, whereas under the arsenite-stress condition only 1 replicate was obtained for 2 transcripts and no data were obtained for 8 transcripts. Transcripts with incomplete stress-condition data were not excluded; the samples that were available were used as they were.
When evaluating the loss, a mask based on the original sequence length was applied so that zero-padded positions did not contribute to the calculation.
Sequence lengths in the training data ranged from 104 to 34,626 nt, with a mean of 2,394.5 nt. The maximum input length was set to 32,768 nt, and sequences exceeding this length were truncated at the 3′ end. Shorter sequences were padded at the 3′ end with the pad token (<pad>) of the NTv3 tokenizer to a length of 32,768 tokens. The corresponding targets (cosine similarity) were padded with zeros, and padded positions were excluded from the loss. Under the unstressed condition, 221,108 of 58,220,439 positions were positive; under the stress condition, 167,402 of 46,558,835 positions were positive.
The TISCA2 cosine similarity profile was used as the continuous target at each position without normalization. Missing values were masked and, together with positions having negative cosine similarity, were excluded from both the loss and the evaluation. For the BCE target and PR-AUC evaluation, positions with cosine similarity ≥ 0.2 were labelled positive.
The data were split into ten folds at the level of transcript identifier. Files derived from the same transcript across the unstressed condition, the stress condition and replicates were assigned to the same fold, so that no transcript was shared between the training and validation sets. Different isoforms of the same gene could be assigned to different folds.

#### IncepTIS Architecture

IncepTIS uses NTv3 (InstaDeepAI/NTv3_100M_pre) as an RNA sequence encoder that extracts a per-nucleotide embedding. NTv3 was selected because it provides single-nucleotide resolution while capturing long-range sequence context.

Let $L$ denote the length of the RNA sequence, $\mathbf{1}_L$ a vector of ones, and [;] concatenation along the feature dimension at each position. Uracil is replaced by thymine in the input sequence, which is passed to NTv3, and a 768-dimensional feature vector is obtained for each nucleotide position from the final layer, denoted $h_{ntv3} \in \mathbb{R}^{\{L\times 768\}}$. The condition is encoded as $c \in \{0, 1\}$,with 0 for the unstressed condition and 1 for the arsenite-stress condition, broadcast across all sequence positions and concatenated with the NTv3 features, giving $h_{in} = [h_{ntv3}; c \cdot \mathbf{1}_L] \in \mathbb{R}^{\{L\times 769\}}$ as the input to the Attention U-Net prediction head.

The Attention U-Net prediction head consists of three downsampling convolutional blocks and three upsampling convolutional blocks, with an attention gate on each of the three skip connections. The model has 108M parameters in total, of which 1.73M belong to the Attention U-Net. The purpose of adding an Attention U-Net on top of the final NTv3 layer is to capture multi-scale sequence features, from global sequence context to local features, and thereby to represent condition-dependent changes in initiation.

The channel widths of the three downsampling blocks are 64, 128 and 256, the bottleneck width is 512, and the channel widths of the three upsampling blocks are 256, 128 and 64. The first convolution used a kernel size of 7 and all other convolutions a kernel size of 3, with a stride of 1 throughout. Max pooling with a kernel size of 2 is applied between downsampling blocks, and each upsampling block uses nearest-neighbour upsampling followed by a convolution with a kernel size of 3. Each convolutional block applies batch normalization, ReLU and dropout with a rate of 0.1.

At each attention gate, the feature from the upsampling path and the corresponding feature from the downsampling path are each projected to an intermediate number of channels by a 1 × 1 convolution followed by batch normalization. The two projections are summed, and ReLU, dropout, a 1 × 1 convolution, batch normalization and a sigmoid are applied in turn to obtain position-wise weights for the skip feature. The weighted skip feature is then added to the feature from the upsampling path.

The final layer of the Attention U-Net head applies a 1 × 1 convolution and predicts one logit $y_i$ per nucleotide $(i = 1, \ldots, L)$ . The regression output is obtained by applying a sigmoid to this logit, $O_{cont,i} = \sigma(y_i)$, and the binary output is obtained from the same logit by subtracting a learnable threshold $\tau$ in logit space, $O_{bin,i} = y_i - \tau$. (Extended Data Fig. 8).

**Loss Function**

The objective of the model is to predict translation initiation sites within a sequence, given the transcript sequence and the presence or absence of stress as input.

The loss was defined as a weighted sum of a binary cross-entropy (BCE) loss and a mean squared error (MSE) loss. Let $V$ denote the set of valid nucleotide positions excluding padding and $|V|$ its cardinality;

$$\mathcal{L} = 0.5\mathcal{L}_{MSE} + 0.5\mathcal{L}_{BCE},$$

$$\mathcal{L}_{MSE} = \frac{1}{|V|}\sum_{i \in V}\left\| CosSim_i - O_{cont,i} \right\|^2,$$

$$\mathcal{L}_{BCE} = -\frac{1}{|V|}\sum_{i \in V}\left( Mask_i \cdot \log \sigma(O_{bin,i}) + (1 - Mask_i) \cdot \log\left(1 - \sigma(O_{bin,i})\right)\right),$$

where $Mask_i$ is 1 at positions whose cosine similarity is greater than or equal to the threshold of 0.2 and 0 otherwise;

$$Mask_i = \begin{cases} 1, & \text{if } CosSim_i \geq 0.2 \\ 0, & \text{Otherwise} \end{cases}.$$

For the BCE loss, $Mask_i$ was used as the target label and BCEWithLogitsLoss was computed on the binary output $O_{bin,i}$. For the MSE loss, the error between the cosine similarity and the regression output was computed over the masked valid positions. The learnable threshold $\tau$ was initialized at logit(0.2), and optimized jointly with the other model parameters.

**Model Training**

Weight decay was set to 0. The batch size was 4 per GPU, giving a global batch size of 16. TF32 was used for matrix multiplications and convolutions. A random seed of 524 was used for both data splitting and training.

Each fold was trained for a total of 69,880 optimizer updates. The learning rate was increased from $1 \times 10^{-6}$ to $1 \times 10^{-3}$ over the first 100 updates and then decayed on a cosine schedule to reach $1 \times 10^{-6}$ at the end of the 20th epoch. The final checkpoint at epoch 20 was used for evaluation in each fold.

Training was performed on four NVIDIA H100 GPUs on TSUBAME 4.0. Training for 20 epochs took 4.9 hours per fold on average. The software environment was Python 3.12.3, PyTorch 2.8.0+cu128 with CUDA runtime 12.8, and timm 1.0.22.

**IncepTIS-evolve Sequence Optimization**

IncepTIS-evolve takes as configuration the total sequence length, the positions of the target initiation sites, and the sequence segments to be held fixed during the search. For Figure 6f–h, each construct contained a 282-nt designed region followed by a fixed GCCACC Kozak sequence and the Renilla luciferase coding sequence. The Renilla luciferase start codon began at nucleotide 289 (1-based numbering). uORFs were specified at positions 61 (uORFx1); 31 and 151 (uORFx2); and 31, 111 and 191 (uORFx3). The downstream luciferase sequence was set to begin at position 289. Of the 1,024 sequences in the initial population, 64 were generated with Evo 2 (evo2_1b_base) and then overwritten at the fixed segments, whereas in the remaining sequences the mutable positions were sampled uniformly from A, C, G and T.

Let $\mu_u^N$ and $\mu_u^S$ denote the mean predicted TISCA2 cosine-similarity score at the specified uORF sites under the unstressed and stress conditions, respectively; $l^N$ and $l^S$ the TISCA2 cosine-similarity scores at the luciferase start site under the two conditions; $d_u$ denotes the differences between unstressed and stress scores at the specified uORF start sites ($d_u = \mu_u^N - \mu_u^S$); and $m_{internal}$ the maximum output under either condition across the region excluding ±4 nt around the target positions. The maximized fitness was defined as

$$F = w_{pair}\, min(\mu_u^N, l^S) + w_{u,N}\mu_u^N + w_{luc,S}l^S + w_{ind}(l^S - l^N) + w_{diff}d_u - w_{luc,N}l^N - w_{u,S}\mu_u^S - w_{int}m_{internal},$$

with $w_{pair} = 10.0$, $w_{u,N} = 1.3$, $w_{luc,S} = 11.5$, $w_{ind} = 11.5$, $w_{diff} = 1.95$, $w_{luc,N} = 4.0$, $w_{u,S} = 2.45$, $w_{int} = 0.5$.

The first term is large only when both $\mu_u^N$ and $l^S$ are high, and therefore suppresses solutions in which only one of the two conditions has been optimized. Each candidate was scored with IncepTIS fold 1, processed in batches of 256 sequences.

In each generation, the 48 sequences with the highest fitness were carried over unchanged to the next generation. The remainder were generated by selecting two parents by tournament selection among 5 candidates, applying uniform crossover in which each mutable position was exchanged with probability 0.5, and then introducing mutations. At mutable positions within 8 nt of a target start codon, each nucleotide was substituted with one of the three alternative bases with probability 0.12; elsewhere the per-nucleotide probability was 0.035. In addition, up to 128 uniformly random sequences were introduced in each generation to maintain the population size. Each of the three designs was run for 600 generations with seed 2772.

**Definition of Success Output**

Success was defined as simultaneously satisfying $\mu_u^N \geq 0.12$, $l^N \leq 0.08$, $l^S \geq 0.08$, $\mu_u^S \leq 0.14$, $l^S - l^N \geq 0.04$, $d_u \geq 0.04$. Among the sequences meeting all of these criteria, the one with the highest fitness was returned; only when no sequence met all criteria was the sequence with the highest fitness among all candidates returned instead. The algorithm is described in Supplementary Algorithms 1 and 2 and Supplementary Tables 1–3.

**Surrogate-Model-Guided Proposal**

Because evaluation by IncepTIS is computationally expensive, it was limited to 1024 sequences per generation. A surrogate model approximating sequence fitness was therefore trained on pairs of evaluated sequences and their fitness values, and used to pre-screen an unevaluated candidate pool at low computational cost. An extremely randomized trees (ExtraTrees) regressor is used as the surrogate model. The model comprised 64 trees (max_features="sqrt", min_samples_leaf=2, bootstrap=True) and was trained from generation 20 once at least 4,096 observations were available, with retraining every generation. When more than 12,000 observations were available, 12,000 were sampled without replacement with probabilities proportional to the fitness of each observation minus the minimum observed fitness, plus $10^{-6}$. Sequences were encoded as 1,096-dimensional A/C/G/T one-hot vectors across 274 mutable positions.

The candidate pool comprised 4096 sequences, assembled after removal of duplicates from a mixture of mutated elites (55%) crossover between elites followed by mutation (30%) and uniformly random sequences (15%). For each candidate c, the mean $\hat{\mu}(c)$and standard deviation $\hat{\sigma}(c)$ of the per-tree predictions $h_b(\psi(c))$ were computed, and the upper confidence bound

$$\alpha(c) = \hat{\mu}(c) + \beta\,\hat{\sigma}(c)$$

was used as the acquisition function（$\beta$= 0.75）. Because the variance between trees becomes large in regions where training data are sparse, it was used as an approximation of predictive uncertainty. The 128 sequences with the highest α were introduced into the next generation, replacing an equal number of random sequences.

### Extended Data Figures

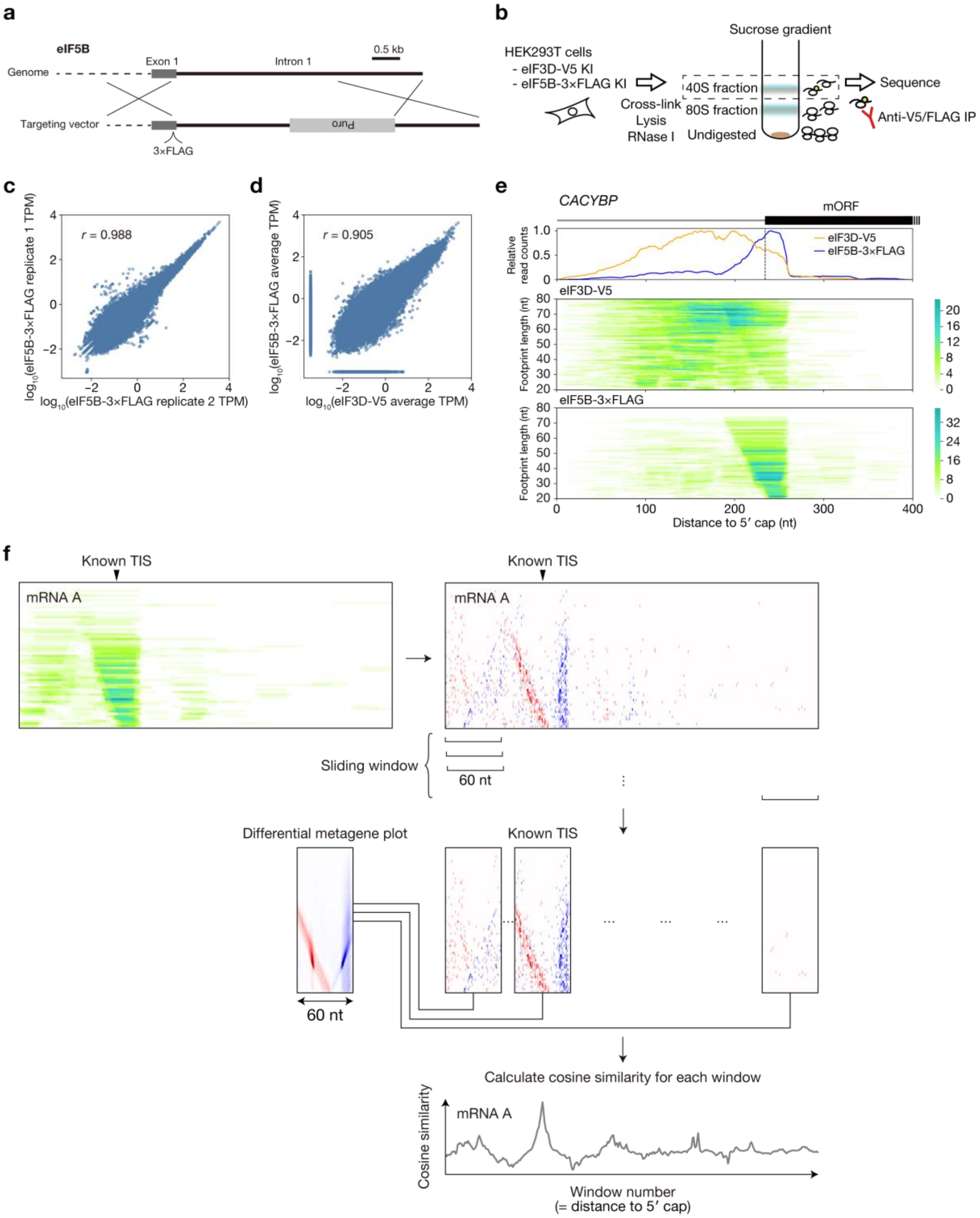


**Extended Data Fig. 1 | Establishment and characterization of eIF5B Sel-TCP-seq.** **a**, Schematic of the strategy used to introduce a 3×FLAG tag at the endogenous *EIF5B* locus in HEK293T cells. The genomic locus and targeting vector are shown. **b**, Schematic of Sel-TCP-seq using eIF3D-V5 or eIF5B-3×FLAG knock-in HEK293T cells. **c**, Reproducibility of eIF5B-3×FLAG Sel-TCP-seq between two biological replicates. Each point represents a transcript. The Pearson's correlation coefficient is shown. **d**, Comparison of transcript-level footprint abundance between eIF3D-V5 and eIF5B-3×FLAG Sel-TCP-seq. Values represent the average of two biological replicates. The Pearson's correlation coefficient is shown. **e**, Representative Sel-TCP-seq profiles at the *CACYBP* transcript. Relative read distributions of eIF3D-V5 and eIF5B-3×FLAG are shown above two-dimensional footprint distributions plotted by footprint length and distance from the 5′ cap. The annotated main ORF (mORF) is indicated. **f**, Schematic of the differential-footprint and cosine-similarity analyses used for TISCA2. Two-dimensional footprint distributions were converted to differential plots by edge detection. Differential footprint distributions were scanned using overlapping 60-nt windows, and cosine similarity between each window and the reference eIF5B differential metagene pattern was calculated to generate a position-resolved TIS similarity score.

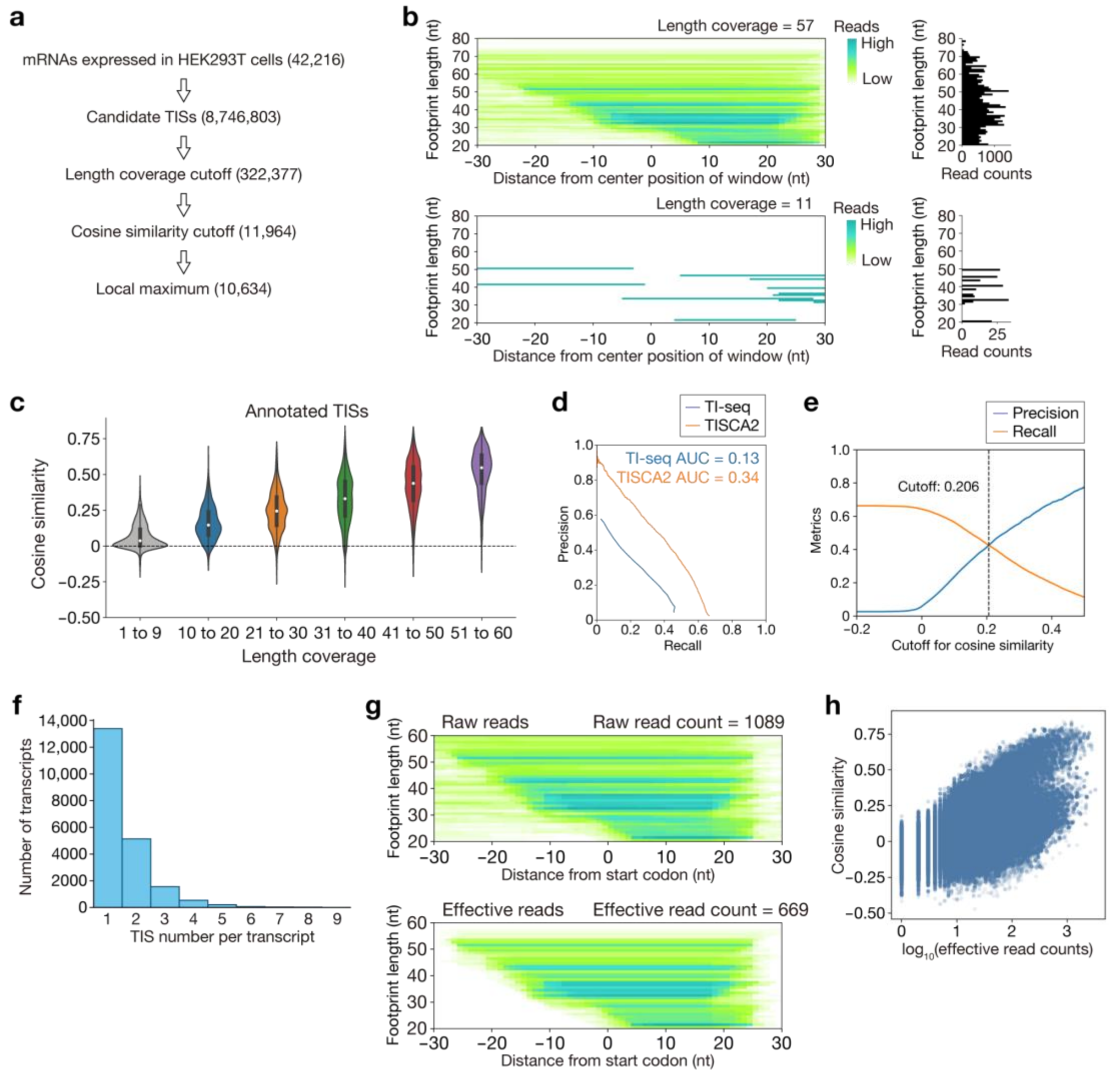


**Extended Data Fig. 2 | Optimization and evaluation of the TISCA2 calling procedure.** **a**, Overview of the TISCA2 TIS-calling workflow and the number of candidate sites retained at each filtering step. **b**, Representative eIF5B footprint distributions with high and low footprint-length coverage. Read-count distributions by footprint length are shown on the right. **c**, Cosine similarity of annotated canonical TISs grouped according to footprint-length coverage. **d**, Precision–recall curves comparing TISCA2 and TI-seq for identification of annotated canonical TISs. Areas under the curve (AUCs) are indicated. **e**, Precision and recall across cosine-similarity cutoffs. The cutoff used for TISCA2 analysis (0.206) is indicated. **f**, Distribution of the number of TISs identified per transcript. **g**, Representative comparison of raw and effective eIF5B footprint reads around an annotated start codon. Effective reads were defined as footprints whose 3′ ends mapped between +15 and +30 nt relative to the start codon. **h**, Relationship between cosine similarity and effective read counts at annotated canonical TISs.

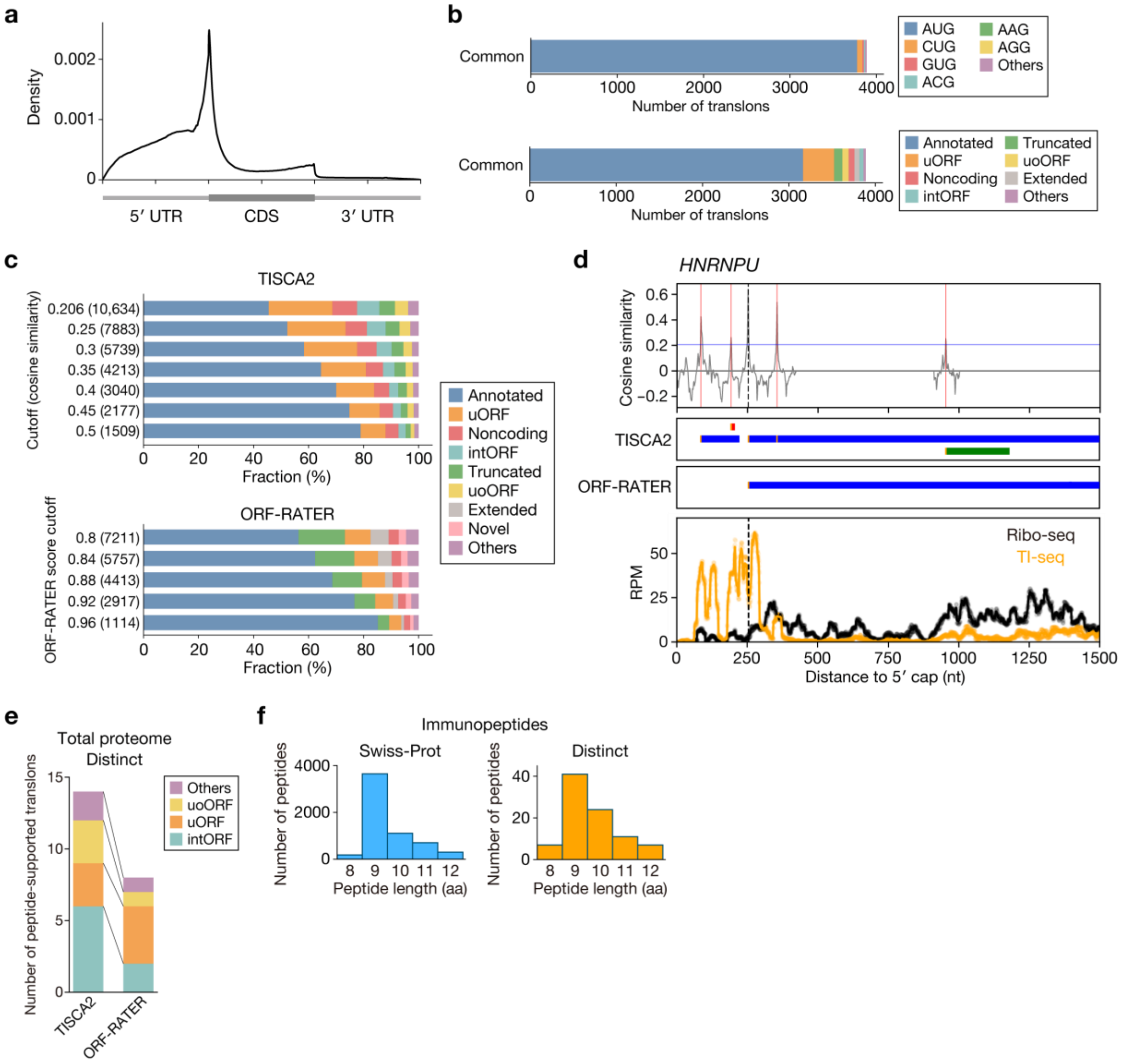

**Extended Data Fig. 3 | Comparison of TISCA2 and ORF-RATER predictions.** **a**, TI-seq metagene profile across 5′ UTRs, CDSs and 3′ UTRs in HEK293T cells. **b**, Start-codon composition and translon classification of the 3,889 translons identified by both TISCA2 and ORF-RATER. **c**, Translon-category distributions obtained across increasing cosine-similarity cutoffs for TISCA2 and ORF-RATER score cutoffs. The number of predicted translons at each cutoff is indicated. **d**, Representative analysis of the *HNRNPU* locus. Cosine similarity, TISCA2 and ORF-RATER predictions, and Ribo-seq and TI-seq profiles are shown. The TISCA2-defined intORF was supported by an N-terminal peptide. **e**, Distinct translons supported by total proteomics using TISCA2- or ORF-RATER-derived search databases, classified according to translon category. **f**, Length distributions of HLA class I immunopeptides assigned to Swiss-Prot proteins and distinct translons.

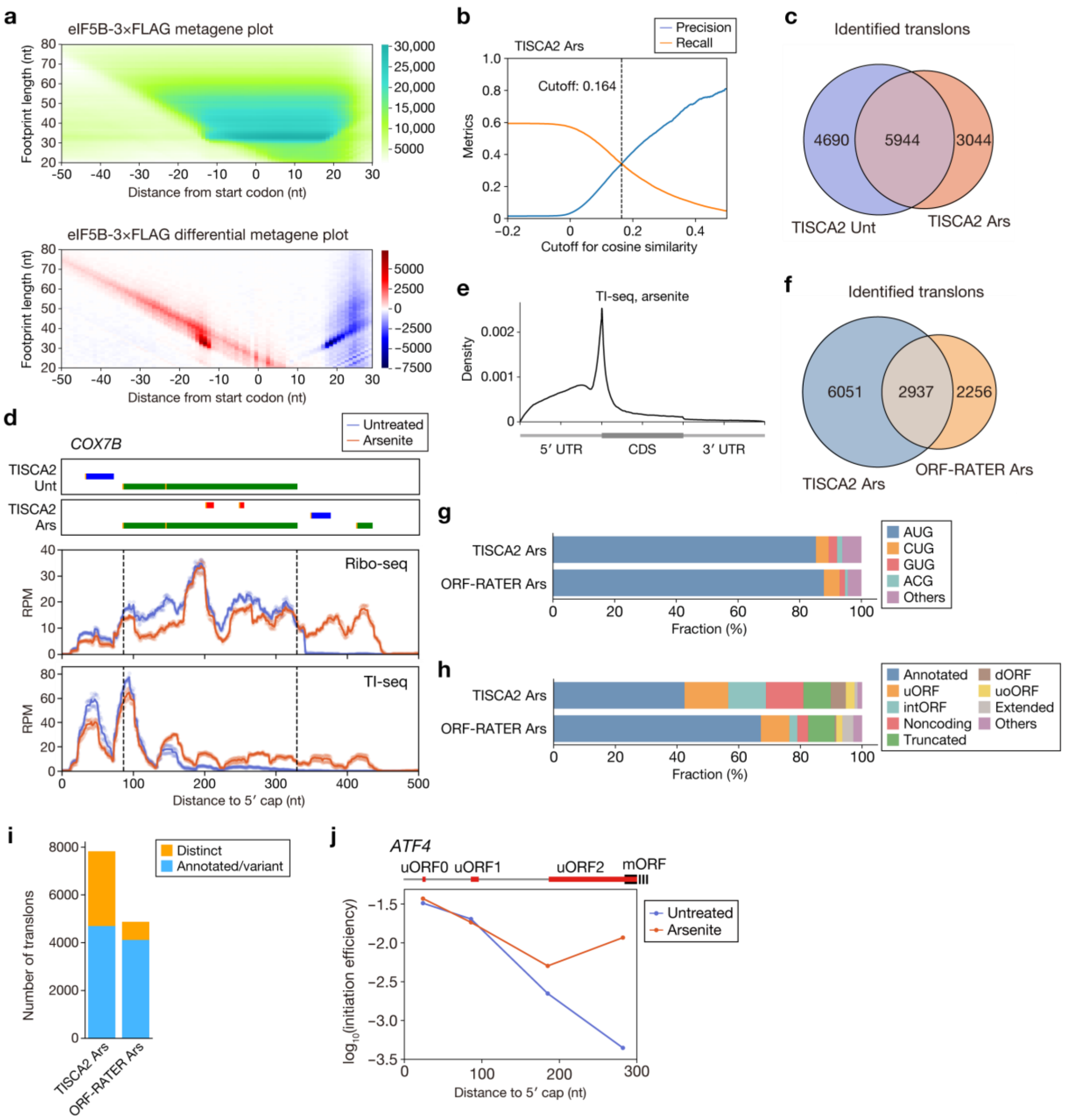


**Extended Data Fig. 4 | Comparison of translation initiation in untreated and arsenite-treated cells.**

**a**, Two-dimensional eIF5B-3×FLAG metagene and differential metagene plots around annotated start codons in arsenite-treated HEK293T cells. **b**, Precision and recall across cosine-similarity cutoffs for TISCA2 analysis of arsenite-treated cells. The cutoff used for TIS calling is indicated. **c**, Overlap between TISs identified by TISCA2 in untreated and arsenite-treated cells. **d**, Representative analysis of the *COX7B* transcript. TISCA2 predictions in untreated and arsenite-treated cells are shown together with Ribo-seq and TI-seq profiles. **e**, TI-seq metagene profile across 5′ UTRs, CDSs and 3′ UTRs in arsenite-treated cells. **f**, Overlap between translons identified by TISCA2 and ORF-RATER after arsenite treatment. **g**, Start-codon composition of translons identified by TISCA2 and ORF-RATER after arsenite treatment. **h**, Translon classification of TISCA2- and ORF-RATER-defined translons after arsenite treatment. **i**, Numbers of annotated/variant and distinct translons identified by TISCA2 and ORF-RATER after arsenite treatment. **j**, IE of TISs within the *ATF4* transcript in untreated and arsenite-treated cells. The positions of uORF0, uORF1, uORF2 and the main ORF (mORF) are indicated.

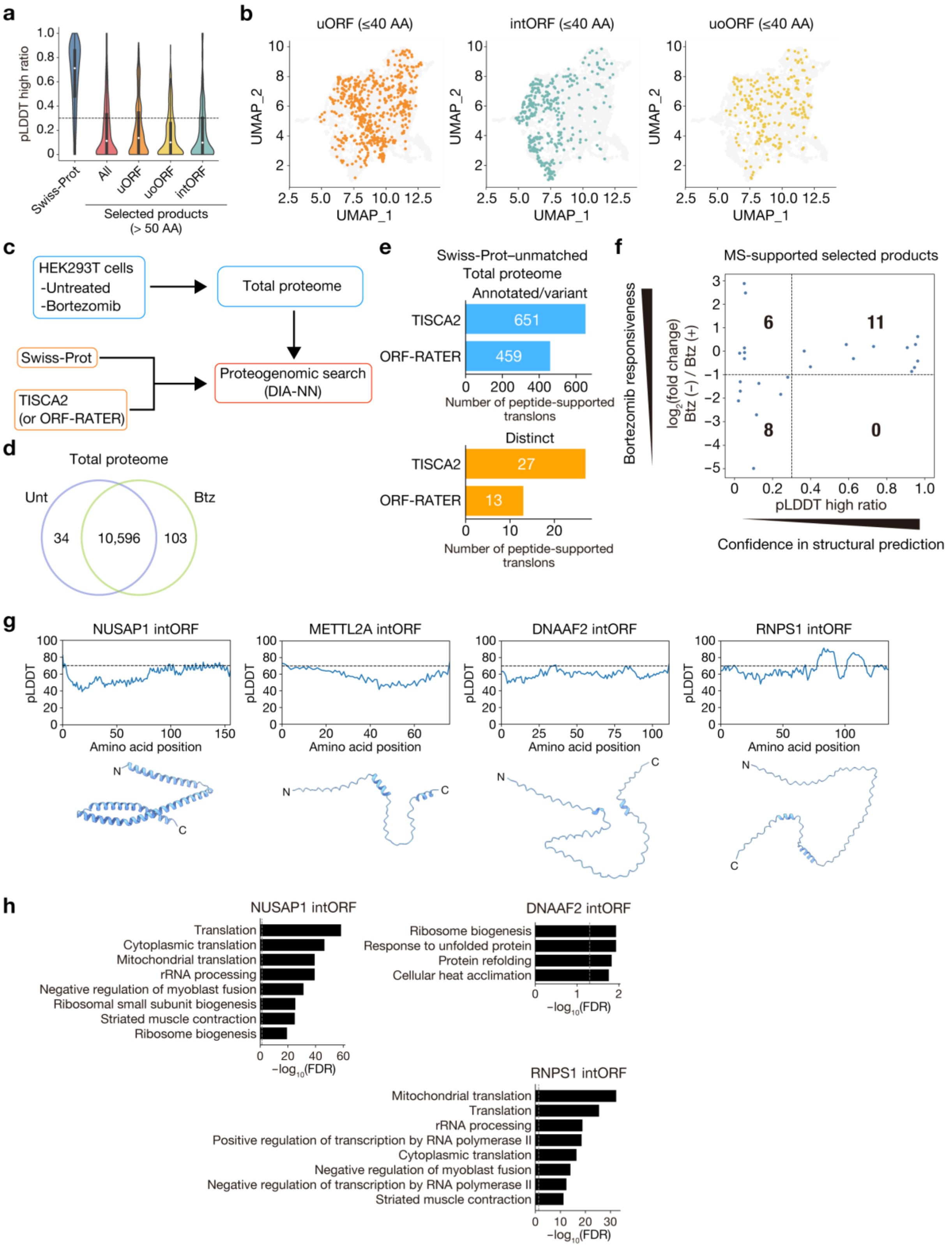


**Extended Data Fig. 5 | Structural prediction and proteomic characterization of TISCA2-predicted translon products.**

**a**, Distribution of pLDDT-high ratios for Swiss-Prot proteins and selected products longer than 50 amino acids, classified by translon category. **b**, Distribution of uORF-, intORF- and uoORF-derived products of 21-40 amino acids in ESM-2/UMAP sequence space. **c**,

Schematic of the total-proteome proteogenomic search. Total proteomes from untreated and Btz-treated HEK293T cells were searched against Swiss-Prot combined with TISCA2- or ORF-RATER–predicted products using DIA-NN. **d**, Overlap of products detected by total proteomics in untreated and Btz-treated cells. **e**, Numbers of Swiss-Prot–unmatched annotated/variant and distinct translons supported by total proteomics using TISCA2- or ORF-RATER–derived search databases. **f**, Relationship between Btz responsiveness and pLDDT-high ratio for MS-supported selected products. Btz responsiveness is shown as the $\log_2$ abundance ratio of untreated to Btz-treated cells. Dashed lines indicate a pLDDT-high ratio of 0.3 and a $\log_2$ abundance ratio of −1. Numbers indicate the number of products in each quadrant. **g**, AlphaFold 3-predicted structures and residue-level pLDDT profiles of the NUSAP1, METTL2A, DNAAF2 and RNPS1 intORF-derived products. Dashed lines indicate pLDDT = 70. N and C indicate the amino and carboxyl termini, respectively. **h**, Gene Ontology enrichment analysis of proteins co-purifying with the NUSAP1, DNAAF2 and RNPS1 intORF-derived products in V5 immunoprecipitation–mass spectrometry experiments. Among the Gene Ontology terms with FDR < 0.05, those with the lowest FDR are shown.

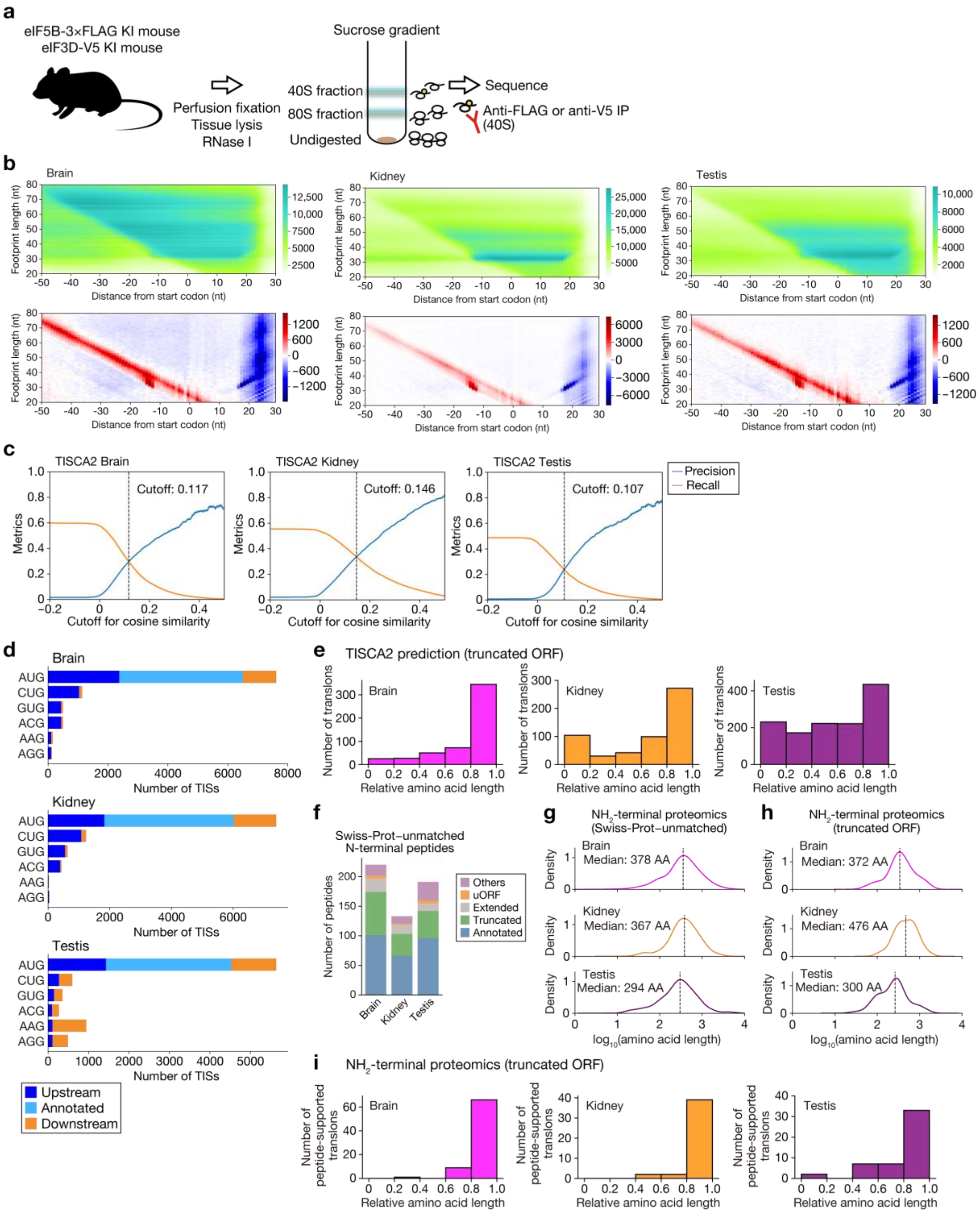

**Extended Data Fig. 6 | Characterization of tissue-specific translation initiation by TISCA2.**

**a**, Schematic of eIF5B-3×FLAG and eIF3D-V5 Sel-TCP-seq in mouse tissues. **b**, Two-dimensional eIF5B-3×FLAG metagene and differential metagene plots around annotated start codons in brain, kidney and testis. **c**, Precision and recall across cosine-similarity cutoffs for TISCA2 analysis of brain, kidney and testis. The cutoffs used for TIS calling are indicated. **d**,

Numbers of TISs initiated at the indicated codons in brain, kidney and testis, classified according to their positions as upstream, annotated or downstream. **e**, Distributions of relative amino-acid lengths of TISCA2-predicted truncated ORFs in brain, kidney and testis. Relative length was calculated against the corresponding annotated product. **f**, Swiss-Prot–unmatched N-terminal peptides detected in brain, kidney and testis, classified according to translon category. **g**, Length distributions of products assigned to Swiss-Prot–unmatched N-terminal peptides in the indicated tissues. Median lengths are indicated. **h**, Length distributions of truncated ORF-derived products supported by N-terminal proteomics. Median lengths are indicated. **i**, Distributions of relative amino-acid lengths of truncated ORF-derived products supported by N-terminal proteomics in brain, kidney and testis. Relative length was calculated against the corresponding annotated product.

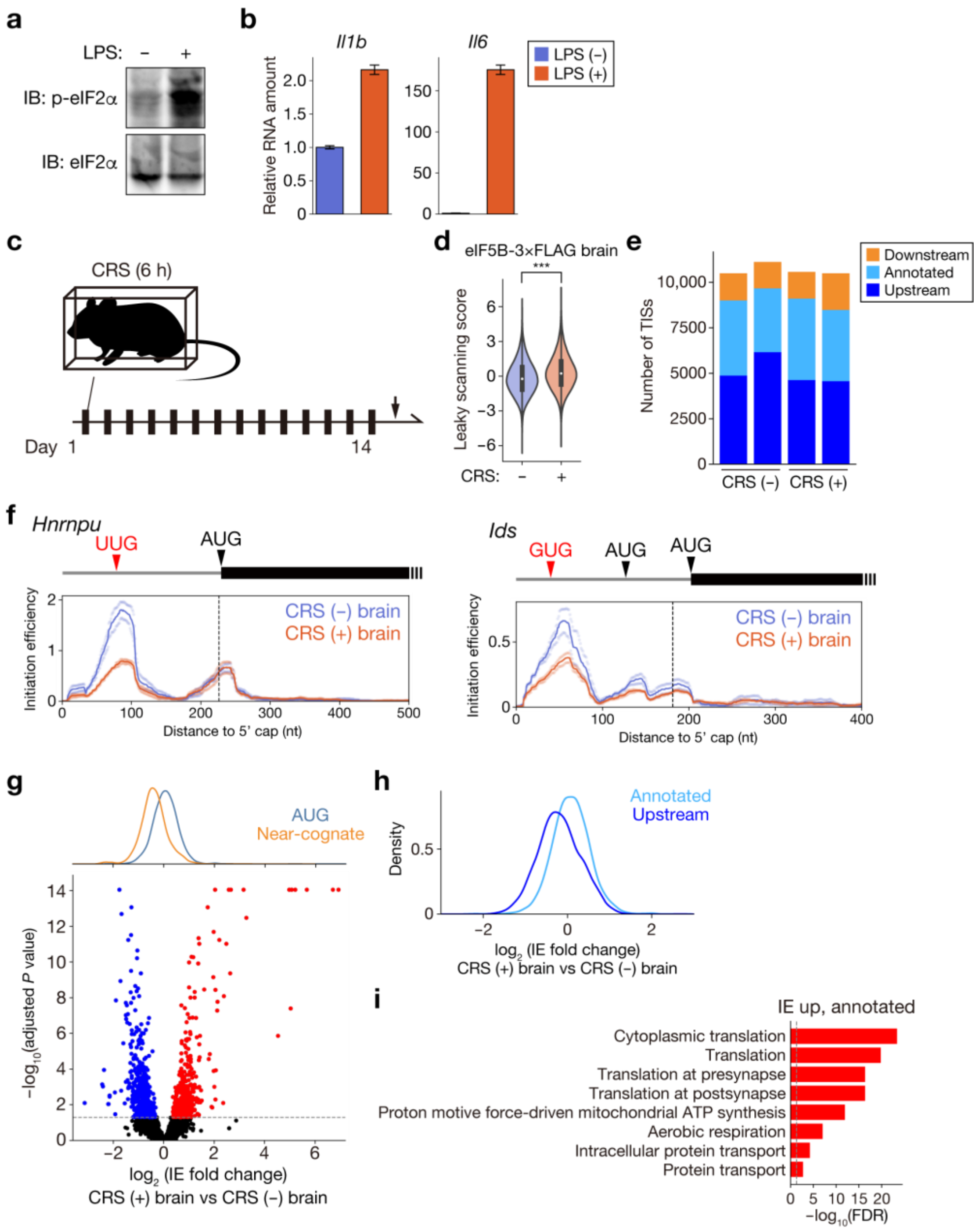


**Extended Data Fig. 7 | Characterization of LPS-treated kidneys and CRS-treated brains.**

**a,** Immunoblot analysis of phosphorylated eIF2α (p-eIF2α) and total eIF2α in kidneys collected 16 h after intravenous administration of LPS (5 mg/kg) or PBS. **b,** Relative *Il6* and *Il1b* mRNA levels measured by quantitative PCR in kidneys collected as in **a**. **c,** Schematic of the chronic restraint stress (CRS) protocol. Mice were subjected to restraint stress for 6 h per day for 14 consecutive days before brain collection. **d,** Leaky-scanning scores calculated from eIF5B-3×FLAG-associated footprints in control and CRS-treated brains. ***$P < 0.001$, Steel-Dwass test. **e,** Numbers of TISCA2-identified TISs located upstream of, at or downstream of annotated CDS start sites in control and CRS-treated brains. **f,** Representative initiation efficiency (IE) profiles of the *Hnrnpu* and *Ids* transcripts in control and CRS-treated brains. The positions and start codons of the indicated TISs are shown. **g,** Genome-wide changes in IE following CRS (*n* = 2 biologically independent samples). AUG- and near-cognate-initiated TISs are indicated separately. Fold changes in IE are plotted against adjusted *P* values, and density distributions are shown above. **h,** Distributions of IE fold changes following CRS for annotated and upstream TISs. **i,** Gene Ontology enrichment analysis of genes showing increased IE at annotated TISs following CRS. Among the Gene Ontology terms with FDR < 0.05, those with the lowest FDR are shown.

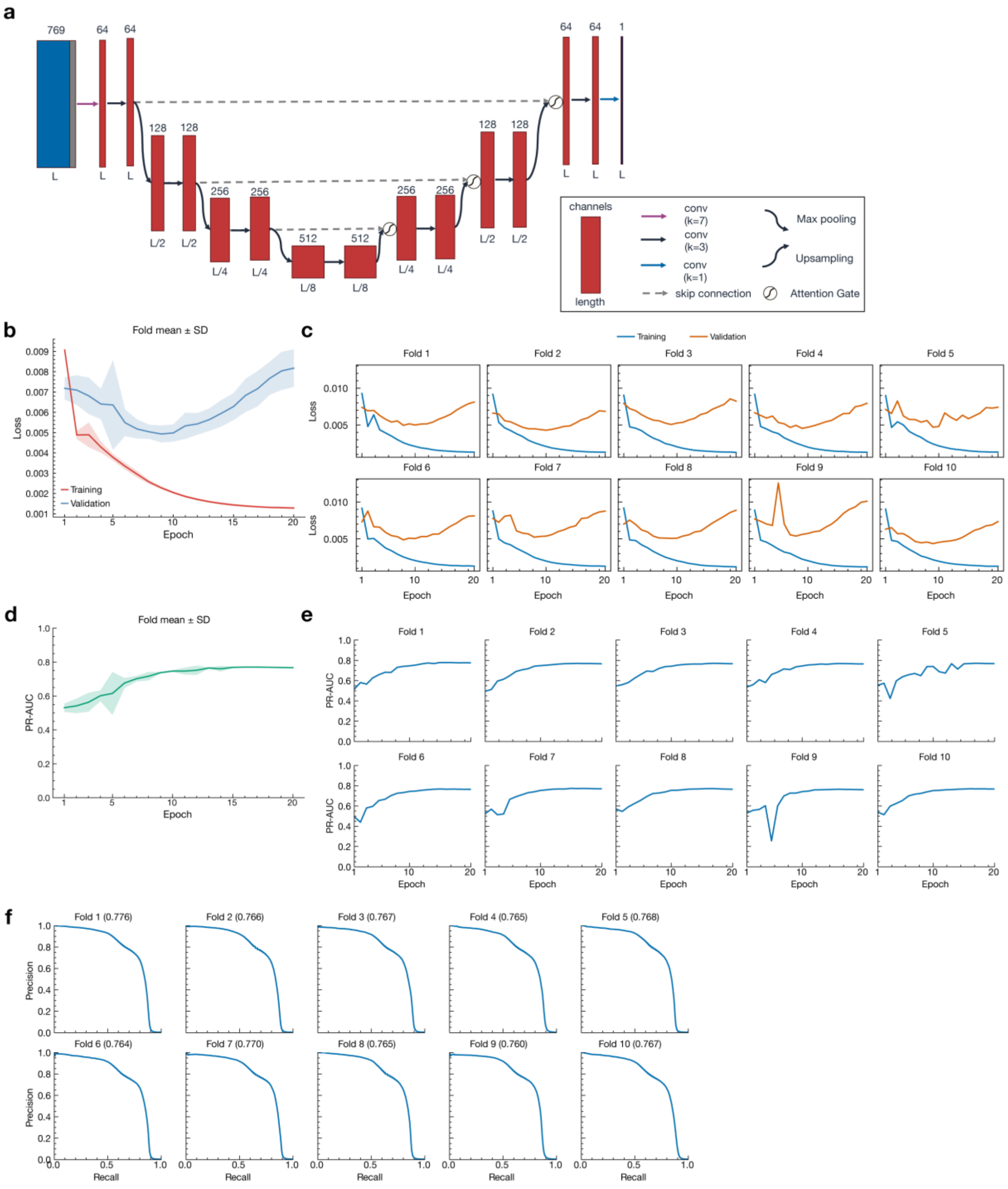


**Extended Data Fig. 8 | Architecture and training performance of IncepTIS.**
**a,** Architecture of the Attention U-Net prediction head. The input comprises 768-dimensional NTv3 sequence representations concatenated with a cellular-state indicator. Numbers above the blocks indicate channel counts, and labels below indicate sequence lengths. L, input sequence length; k, convolutional kernel size. Skip connections and attention gates are indicated. **b,c,** Training and validation loss across 20 epochs, summarized as the mean ± s.d. across ten cross-validation folds (**b**) and shown separately for each fold (**c**). **d,e,** Validation precision–recall area under the curve (PR-AUC) across training epochs, summarized as the

mean ± s.d. across ten folds (**d**) and shown separately for each fold (**e**). **f,** Precision–recall curves for held-out predictions in each cross-validation fold. PR-AUC values are indicated in parentheses. Nucleotide positions with measured TISCA2 cosine similarity ≥0.2 were defined as positive. Cross-validation splits were assigned by transcript identifier, with all conditions and replicates of each transcript retained in the same fold.

**Supplementary Table**

**Supplementary Table 1** IncepTIS-evolve Parameters (see also Supplementary Algorithms 1 and 2).

| Symbol | Definition |
|---|---|
| $D$ | Design specification, $D = (L, \mathcal{F}, S)$. |
| $L$ | Length of the designed sequence (nt). |
| $\mathcal{F}$ | Set of fixed segments, each given by a start position and a sequence; not altered during the search. |
| $S$ | Set of target sites (the uORFs and the luciferase start). |
| $\mathcal{M}$ | Set of mutable positions, i.e. positions not covered by $\mathcal{F}$. |
| $m$ | Number of mutable positions, $m = \lvert\mathcal{M}\rvert$. |
| $C$ | Set of hard constraints used for the success criterion. |
| $\Sigma$ | Nucleotide alphabet, $\Sigma = \{A, C, G, T\}$. |
| $w$ | Vector of objective weights, $w = (w_{pair}, w_{u,N}, w_{luc,S}, w_{ind}, w_{diff}, w_{luc,N}, w_{u,S}, w_{int})$; individual entries are defined in supplementary methods. |
| $P$ | Population size. |
| $G$ | Total number of generations. |
| $E$ | Number of elite individuals carried over unchanged. |
| $k$ | Tournament size. |
| $R$ | Number of random immigrants introduced per generation. |
| $p_{mut}$ | Per-base mutation probability outside the target neighbourhood. |
| $p_{loc}$ | Per-base mutation probability within the target neighbourhood. |
| $g_0$ | Generation at which surrogate-guided proposal starts. |
| $\Delta$ | Refitting interval of the surrogate model (generations). |
| $N_{min}$ | Minimum number of observations required to fit the surrogate model. |
| $M$ | Size of the candidate pool, $M = \lvert\mathcal{P}\rvert$. |
| $n$ | Number of proposals introduced per generation, $n \leq R$. |
| $B$ | Number of trees in the Extra Trees ensemble. |
| $\beta$ | Exploration coefficient, $\beta \geq 0$. |

**Supplementary Table 2** IncepTIS-evolve Variables (see also Supplementary Algorithms 1 and 2).

| Symbol | Definition |
|---|---|
| $x^*$ | Final output sequence; $x_c$ if $x_c \neq \perp$, otherwise $x_b$. |
| $X$ | Population of the current generation, $\lvert X \rvert = P$. |
| $\mathcal{H}$ | Observation history, $\mathcal{H} = \{(x_j, F(x_j))\}$. |
| $x_b$ | Sequence with the highest fitness, irrespective of the constraints. |
| $x_c$ | Sequence with the highest fitness among those satisfying all constraints. |
| $\perp$ | Undefined; no such sequence has been found. |
| $g$ | Generation index, $g = 0, \dots, G$. |
| $x$ | A sequence, $x \in \Sigma^L$. |
| $a^N$ | Position-wise TIS activity profile under the untreated condition. |
| $a^S$ | Position-wise TIS activity profile under the arsenite stress condition. |
| $\varphi(x)$ | Summary metrics aggregated from the profiles. |
| $F(x)$ | Fitness of sequence $x$. |
| $\pi$ | Permutation of indices sorting $F$ in descending order; $x_{\pi(j)}$ is the $j$-th ranked sequence. |
| $j$ | Rank index. |
| $\mathcal{E}$ | Set of elite sequences, $\lvert \mathcal{E} \rvert = E$. |
| $X'$ | Population of the next generation under construction. |
| $p_1, p_2$ | Parent sequences chosen by tournament selection. |
| $\Pi$ | Set of proposals returned by ProposeBO, $\lvert \Pi \rvert \leq n$. |
| $\mathcal{T}$ | Training set drawn from $\mathcal{H}$ for the surrogate model. |
| $b$ | Tree index, $b = 1, \dots, B$. |
| $h_b$ | Prediction function of tree $b$, $\{0,1\}^{4m} \to \mathbb{R}$. |
| $\mathcal{P}$ | Pool of unevaluated candidate sequences. |
| $c$ | A candidate sequence, $c \in \mathcal{P}$. |
| $\hat{\mu}(c)$ | Mean prediction across the trees. |
| $\hat{\sigma}(c)$ | Standard deviation of the predictions across the trees. |
| $\alpha(c)$ | Acquisition function. |

**Supplementary Table 3** Functions and Operators in IncepTIS-evolve (see also Supplementary Algorithms 1 and 2).

| Symbol | Definition |
|---|---|
| $f_{TIS}$ | The trained IncepTIS model; maps a sequence and a condition to a position-wise TIS activity profile. |
| $\psi$ | One-hot encoding of the mutable positions, $\psi: \Sigma^L \rightarrow \{0,1\}^{4m}$. |
| $\text{argmax}^{(n)}$ | Operator returning the top $n$ elements. |
| InitPopulation | Builds the initial population from Evo 2 generations overlaid with the fixed segments, filling the remaining slots with random sequences. |
| Summarize | Aggregates the position-wise profiles into the summary metrics. |
| Objective | Computes the fitness from the summary metrics and the weights. |
| ArgSortDesc | Sorts the indices in descending order of fitness. |
| Feasible | Tests whether all hard constraints in $C$ are satisfied. |
| Tournament | Selects one parent as the fittest of $k$ randomly drawn candidates. |
| Crossover | Uniform crossover exchanging each mutable position with probability 0.5. |
| Mutate | Substitutes mutable positions with one of the three other nucleotides at probability $p_{mut}$ or $p_{loc}$. |
| ProposeBO | Returns the proposal set obtained from the surrogate model and the acquisition function (Supplementary Algorithm 2). |
| RandomSeq | Generates a random sequence that preserves the fixed positions. |
| Subsample | Draws a training set from $\mathcal{H}$ without replacement, with probability proportional to fitness. |
| FitExtraTrees | Fits an Extra Trees regressor and returns the $B$ trees $h_b$. |
| CandidatePool | Constructs the candidate pool from mutated elites, elite crossovers, and random sequences. |

**Supplementary Algorithm 1** IncepTIS-evolve.

**Require**: design D; weights $w$; constraints C; P; G; E, k, R $\in \mathbb{N}$; $p_{mut}, p_{loc} \in [0,1]$; $\beta \in \mathbb{R}_{\geq 0}, g_0, \Delta, N_{min}$
**Ensure**: sequence $x^* \in \Sigma^L, \Sigma = \{\mathrm{A, C, G, T}\}$

$X \leftarrow$ InitPopulation(D, P)
$\mathcal{H} \leftarrow \emptyset; x_b \leftarrow \perp; x_c \leftarrow \perp$
**for** $g \leftarrow 0$ **to** $G$ **do**
**for all** $x \in X$ **do**
$(a^N, a^s) \leftarrow f_{TIS}(x)$
$\varphi(x) \leftarrow$ Summarize$(a^N, a^s, D)$
$F(x) \leftarrow$ Objective$(\varphi(x), w)$
**end for**
$\mathcal{H} \leftarrow \mathcal{H} \cup \{(x, F(x)): x \in X\}$
$\pi \leftarrow$ ArgSortDesc$(F)$
**if** $x_b = \perp$ **or** $F(x_{\pi(1)}) > F(x_b)$ **then** $x_b \leftarrow x_{\pi(1)}$ **end if**
**for** $j \leftarrow 1$ **to** $|X|$ **do**
**if** Feasible$(\varphi(x_{\pi(j)}), C)$ **then**
**if** $x_c = \perp$ *or* $F(x_{\pi(j)}) > F(x_c)$ **then** $x_c \leftarrow x_{\pi(j)}$ **end if**
**break**
**end if**
**end for**
**if** $g = G$ **then break end if**
$X' \leftarrow \{x_{\pi(1)}, \ldots, x_{\pi(E)}\}$
**while** $|X'| < P - R$ **do**
$(p_1, p_2) \leftarrow$ (Tournament$(X, F, k)$, Tournament$(X, F, k)$)
$X' \leftarrow X' \cup$ {Mutate(Crossover$(p_1, p_2), p_{mut}, p_{loc}$)}
**end while**
**if** $g \geq g_0$ **and** $g \bmod \Delta = 0$ **and** $|\mathcal{H}| \geq N_{min}$ **then**
$X' \leftarrow X' \cup$ ProposeBO$(\{x_{\pi(1)}, \ldots, x_{\pi(E)}\}, \mathcal{H})$
**end if**
**while** $|X'| < P$ **do** $X' \leftarrow X' \cup$ {RandomSeq$(D)$} **end while**
$X \leftarrow X'$
**end for**
**if** $x_c \neq \perp$ **then return** $x_c$ **else return** $x_b$ **end if**

**Supplementary Algorithm 2** ProposeBO.

**Require**: $\mathcal{H}, \mathcal{E}, M, B \in \mathbb{N}; \beta \in \mathbb{R}_{\geq 0}, n$
**Ensure**: proposal set $\Pi$, $|\Pi| \leq n$

```
T ← Subsample(H)
{h_b}_{b=1}^B ← FitExtraTrees({(ψ(x), y): (x, y) ∈ T})
P ← CandidatePool(E, M)
for all c ∈ P do
    μ̂(c) ← B^{-1} Σ_{b=1}^B h_b(ψ(c))
    σ̂(c) ← (B^{-1} Σ_{b=1}^B (h_b(ψ(c)) − μ̂(c))^2)^{1/2}
    α(c) ← μ̂(c) + β σ̂(c)
end for
return argmax^{(n)}_{c∈P\E} α(c)
```

In math notation:

$\mathcal{T} \leftarrow \text{Subsample}(\mathcal{H})$
$\{h_b\}_{b=1}^{B} \leftarrow \text{FitExtraTrees}(\{(\psi(x), y) : (x, y) \in \mathcal{T}\})$
$\mathcal{P} \leftarrow \text{CandidatePool}(\mathcal{E}, M)$
**for all** $c \in \mathcal{P}$ **do**
$\quad \hat{\mu}(c) \leftarrow B^{-1} \sum_{b=1}^{B} h_b(\psi(c))$
$\quad \hat{\sigma}(c) \leftarrow \left(B^{-1} \sum_{b=1}^{B} \left(h_b(\psi(c)) - \hat{\mu}(c)\right)^2\right)^{1/2}$
$\quad \alpha(c) \leftarrow \hat{\mu}(c) + \beta \hat{\sigma}(c)$
**end for**
**return** $\text{argmax}^{(n)}_{c \in \mathcal{P} \setminus \mathcal{E}} \alpha(c)$
